# ACSS2-Mediated Metabolic-Epigenetic Crosstalk Drives Fulvestrant Resistance and Represents a Novel Therapeutic Target

**DOI:** 10.1101/2025.09.23.678089

**Authors:** Ayça N Mogol, Jin Young Yoo, Alicia Arredondo Eve, Mahima Goel, David J Dutton, Claire P Schane, Audrey Lam, Debapriya Dutta, Betsy Barnick, Eda D. Erdogan, Erik R. Nelson, Maria Grosse-Perdekamp, Zeynep Madak Erdogan

**Affiliations:** Division of Nutritional Sciences, University of Illinois, Urbana-Champaign, Urbana, IL, 61801, USA; Cancer Center at Illinois, Urbana, IL, 61801, USA; Department of Food Science and Human Nutrition, University of Illinois, Urbana-Champaign, Urbana, IL, 61801, USA; Carle Illinois College of Medicine, University of Illinois Urbana-Champaign, Urbana, IL, 61801, USA; Department of Molecular and Integrative Physiology, University of Illinois, Urbana-Champaign, Urbana, IL, 61801, USA; Carle Foundation Hospital, Urbana, IL, 61801, USA; Carl R. Woese Institute for Genomic Biology, University of Illinois, Urbana-Champaign, Urbana, IL, 61801, USA

**Author notes:** Corresponding author: Zeynep Madak-Erdogan, Department of Food Science and Human Nutrition University of Illinois, Urbana-Champaign, 1201 W Gregory Dr, Urbana, IL, 61801, USA.

**Keywords:** ACSS2, ER⍰, H3K27ac, epigenetics, therapy resistance, breast cancer, liver metastasis

## Abstract

Endocrine therapies target hormone-dependent cancer cells, primarily through estrogen receptor alpha (ERα), expressed in ∼70% of breast cancers (ER+). Despite treatment advances, 30-40% of ER+ breast cancer patients experience recurrence and metastasis, with 5-year survival rates of only 31.9%. We validated poor outcomes for liver metastasis patients treated with Fulvestrant (Fulv) using the local Carle Foundation Hospital cohort and examined metabolic pathways in liver metastatic patient-derived xenograft (PDX) models, revealing upregulated lipid and acetyl-CoA production. Our previous work demonstrated that combining Fulv with acetyl-CoA synthase inhibitor (ACSI) targeting Acyl-CoA Synthetase Short Chain Family Member 2 (ACSS2), synergistically reduced ER+ metastatic breast cancer (MBC) cell viability in vitro. Using multiple analytical approaches-isotope tracing, CUT&RUN sequencing, immunofluorescence, western blot, and RNA sequencing-we characterized the effects of acetyl-CoA synthesis inhibition on Fulv-induced alterations. Fulv treatment of MBC cells increased ACSS2 expression and acetate utilization. Isotope tracing revealed that Fulv decreased acetate flux to the TCA cycle while promoting fatty acid synthesis. Importantly, ACSS2 was predominantly nuclear and CUT&RUN sequencing showed that Fulv treatment increased ACSS2 chromatin occupancy and ERα/ACSS2/H3K27ac overlapping sites near genes associated with tumor progression, which was eliminated by combination of ACSI and Fulv. RNA sequencing revealed reduction of Fulv-induced expression of genes involved in cancer cell metabolism and key signaling pathways in cancer with the Fulv+ACSI combination. In a therapy-resistant xenograft model, combining Fulv and ACSI reduced Fulv-dependent increase in metastatic burden. Our findings indicate ACSS2 contributes to endocrine therapy resistance through nuclear acetyl-CoA provision for epigenetic alterations. Targeting these cancer cell adaptations represents a novel therapeutic approach potentially reducing metastasis-related mortality and improving breast cancer treatment outcomes.

## 2. Introduction

Breast cancer is the most common cancer diagnosis for women in the US, accounting for 32% of all new cases and serving as the second leading cause of cancer-related deaths at 14% ^1^. The disease is classified into four main subtypes based on estrogen receptor (ER) and human epidermal growth factor receptor 2 (HER2) expression, with the ER+/HER2-subtype representing approximately 70% of cases and having the highest incidence across all age groups ^2^. Distant metastasis is the primary driver of mortality in breast cancer ^3,4^, with five-year survival rates of 31.9% ^5^. More than 75% of the patients express ERα, making the tumors vulnerable to endocrine therapies including aromatase inhibitors, selective estrogen receptor modulators (SERMs), selective estrogen receptor degraders (SERDs) like Fulvestrant (Fulv), and cyclin-dependent kinase 4/6 inhibitors ^2,6,7^.

Unlike other endocrine therapies that compete with estrogen binding, Fulv works by forming a complex with ER that triggers its proteasomal degradation, effectively eliminating the receptor from cancer cells ^6^. However, despite generally better survival rates in ER+ breast cancers ^8^, endocrine therapy resistance remains a critical challenge, affecting patients through a variety of mechanisms, including epigenetic reprogramming, metabolic reprogramming, and activation of alternative growth pathways ^7,9–11^. This resistance is especially problematic in liver metastasis, where the metabolically active hepatic microenvironment may facilitate metabolic adaptations that promote therapy resistance, leading to significantly worse progression-free survival in Fulv-treated patients compared to other metastatic sites ^12,13^. Current approaches to overcome endocrine resistance, including combination therapies with CDK4/6 inhibitors, often provide only temporary benefits, highlighting the urgent need for novel strategies targeting the fundamental metabolic vulnerabilities that drive resistance mechanisms.

Emerging evidence suggests that cancer cells undergo metabolic reprogramming to survive endocrine therapy, with altered acetyl-CoA metabolism playing a central role in both metabolic adaptation and epigenetic regulation of resistance genes ^14,15^. Acyl-CoA Synthetase Short Chain Family Member 2 (ACSS2), an enzyme responsible for synthesizing acetyl-CoA from acetate, is located in both the cytosol and nucleus and has been overexpressed in human breast, ovarian, and lung tumor samples ^16,17^. In the nucleus, ACSS2-derived acetyl-CoA serves as a substrate for histone acetyltransferases, directly linking cellular metabolism to chromatin remodeling and gene expression changes ^18^ that can drive cancer cell survival ^19^ and therapy resistance ^20^. Under stressful conditions, ACSS2 promotes cancer cell survival, as demonstrated in ER+ breast cancer cells where 4-hydroxy tamoxifen treatment upregulated ACSS2 expression and enhanced cell viability (18). We previously demonstrated that ACSS2 expression increases with Fulv treatment in vivo, and inhibition of this enzyme synergizes with Fulv to decrease cell viability in vitro ^15^. However, the mechanistic basis for this synergy, particularly the role of nuclear ACSS2 in epigenetic regulation of endocrine resistance genes, remains unclear.

In this study, we hypothesized that Fulv treatment increases nuclear ACSS2 levels to enhance local acetyl-CoA production, driving histone acetylation and transcriptional activation of genes critical for endocrine therapy resistance. Using RNA sequencing, Cleavage Under Targets & Release Using Nuclease (CUT&RUN) sequencing, isotope tracing assays, and functional studies in patient-derived xenograft (PDX) models, we demonstrate that nuclear ACSS2 plays a crucial role in Fulv-induced epigenetic reprogramming of metastatic breast cancer cells. Our findings reveal that targeting this metabolic-epigenetic axis through ACSS2 inhibition represents a promising strategy for overcoming endocrine therapy resistance. This novel combination approach might provide new strategies to improve outcomes for patients with ER+ metastatic breast cancer.

## 3. Methods

### Retrospective Chart Review

This retrospective study was determined to be Exempt Human Subjects Research by the Institutional Review Board (IRB) at Carle Foundation Hospital, and all methods were carried out in accordance with Carle Foundation Hospital guidelines and regulations (IRB protocol number 21CCC3480). A retrospective chart review was conducted to obtain de-identified information at the Carle Cancer Institute for patients who were diagnosed with metastatic hormone receptor-positive breast cancer from 1/1/2010 to 10/31/2021. Following inclusion criteria was used: 1) Female or male patients with Stage IV receptor-positive (ER/PR +, HER2 -) breast cancer, 2) Clear documentation of a malignancy other than breast cancer in stages 0-III, and final analysis included 192 patients who met all these inclusion criteria. Electronic medical records of the study subjects were reviewed to collect information on 1) diagnosis of metastatic disease, 2) time when treatment for metastasis starts, if different from diagnosis of metastatic disease, type of treatment and medications, 3) information on progression of disease and subsequent line(s) of treatment, 4) information on response to treatment, and 5) patient medical history and general demographics.

Specific retrospective lab values were collected at the time of diagnosis of metastasis and at the time of progression of disease/change of treatment. Patient history and demographics variables (such as age at diagnosis of metastasis, sex, weight, height and BMI at the time of diagnosis of metastasis), metastasis diagnosis date, medication start and end dates (relative to the date of diagnosis of metastasis), family history of cancer or chronic diseases, alcohol/smoking history, menopause status at the time of diagnosis and time of metastasis, metastasis date, metastasis location (liver, bones, lymph nodes, respiratory, other), ER, PR, HER2 status of breast cancer tissue, patient race, survival data, and history of comorbidities such as diabetes and dyslipidemia, were also collected.

Cancer-specific treatment data was collected and divided into 6 categories based on treatment type: hormonal therapy (Tamoxifen, Aromatase inhibitors such as Letrozole, Anastrozole, Exemestane, Fulv) and CDK4/6 inhibitors such as Palbociclib, Ribociclib, Abemaciclib), chemotherapy (Everolimus, Capecitabine, Gemcitabine, Paclitaxel, Docetaxel, Adriamycin, Cyclophosphamide, Vinorelbine, Eribulin, Carboplatin, Abraxane, Sacituzumab), Her-2 directed treatments (Trastuzumab, Ado-trastuzumab emtansine, Enhertu), PARP inhibitors (Olaparib), immunotherapy (Pembrolizumab), and molecular-targeted therapy (Alpelisib). This data was collected at multiple time points: 1) treatment before diagnosis of metastasis, 2) 1st line treatment, 3) 2nd line treatment, 4) 3rd line treatment, 5) 4th line treatment.

### Cell lines

All cell lines were obtained from the American Type Culture Collection (ATCC, Manassas, VA). MCF7 ESR1^D538G^ and MCF7 ESR1^Y537S^ cells that we previously characterized ^14,15,21^. These cells, derived from the MCF7 cell line (RRID:CVCL_0031), carry mutations in the estrogen receptor gene (ESR1) that are exclusively found in patients with metastatic breast cancer and are associated with endocrine therapy resistance. The cells were cultured in Dulbecco’s Modified Eagle Medium (DMEM) with nonessential amino acid (NEAA) salts, 10% fetal bovine serum (FBS), 100 µg/mL penicillin/streptomycin, and 50 mg/mL gentamicin unless otherwise stated.

### Cell line xenograft study

To investigate the impact of the pharmacological ACSS2 inhibition on Fulv-dependent increase in metastatic burden, 1x10^6^ MCF7-ESR1^Y537S^ cells were injected into 5-week-old immunodeficient female NSG (NOD.Cg-*Prkdc^scid^ Il2rg^tm1Wjl^*/SzJ) mice (Jackson Laboratory, stock no. 005557, Bar Harbor, ME, USA). Once a week, the tumors were imaged using the Perkin Elmer IVIS imaging system (RRID:SCR_018621) at the Beckman Institute for Advanced Science and Technology at the University of Illinois Urbana Champaign (UIUC), and twice weekly, mice and food weights were measured. For imaging, mice were injected with 150 mg/kg luciferin (Regis Technologies, Morton Grove, IL; NC0520380) (dissolved in DPBS). 4 weeks later, after tumors were detectable by IVIS imaging in majority of animals, animals were randomized into 4 treatment groups (N=8): Veh, Fulv, ACSS2 inhibitor (ACSI), and Fulv+ACSI Combination. Fulv (Selleckchem; Houston, TX, USA; S1191) (dissolved in 10% DMSO 90% corn oil) treatment was administered by 25 mg/kg/day subcutaneous injections for 5 days a week. Non-Fulv groups received 10% DMSO and 90% corn oil solution in the same volumes by the same methods. On Day 35, ACSI (VY-3-135; Selleckchem; Houston, TX, USA; E1147) (dissolved in 5% DMSO 95% corn oil) treatment started by administering 100 mg/kg/day for 5 days a week by force feeding, based on the previous studies ^22^. Non-ACSI treatment groups received 5% DMSO 95% corn oil solution in the same volumes by the same method. During the study two animals were lost from both the Veh and ACSI groups each and one from the combination, reducing the mice number to 6 for Veh and ACSI groups and to 7 for combination group. After 3 weeks of treatment, all mice are sacrificed, and tumors were collected.

### Patient-Derived Xenograft Experiments

The samples were obtained from the Huntsman Cancer Institute at the University of Utah. The samples were prepared according to the previously described protocols ^23^. Briefly, PDX tumors, HCI044EI and HCI013, were generated with 5–6-week-old, female, NRG mice from JAX/PRR. The fresh fragments of tumors were implanted into the cleared inguinal mammary fat pads (n=10 for each cell line). The mice with fresh fragments had 2 treatment groups: Vehicle (Veh) and Fulv. For Fulv treatment, 100 mg/kg/mice Fulv (Selleckchem; Houston, TX, USA; S1191) (dissolved in 10% DMSO 90% corn oil) was used twice weekly. The mice were terminated 8 weeks after the tumor implantation and tumors were collected.

### Gene expression analysis

#### NanoString metabolic panel

5 Veh and 5 Fulv PDX tumor samples were processed as follows: approximately 20 mg of tumor sample was homogenized with 1 mL of TRIzol reagent (Life Technologies, Carlsbad, CA< USA) for 3 min. After homogenization, the samples were mixed with 250 µL of chloroform and centrifuged. Then, we proceeded to take the clear phase to a new tube and mixed it with 0.5 mL of isopropanol. After overnight incubation at 4 °C, we centrifuged the sample, washed the RNA pellet with 70% EtOH, dried the sample, and resuspended the RNA in DEPC-treated water. We then used the RNeasy mini kit (Qiagen, Hilden, Germany, cat # 74104) to clean up the RNA. After validation of RNA quantity and purity, samples were submitted to the Cancer Center at Illinois, Tumor Engineering and Phenotyping Shared Resource facility for Nanostring assay for the XT-CSO-HMP1-12-Human metabolic panel.

#### Bulk RNA Sequencing

Each experiment consisted of four treatment groups: Veh, Fulv, ACSS2 inhibitor (ACSI), and Fulv+ACSI, or, Veh, Fulv, siACSS2 and siACSS2+Fulv, with three replicates per group. 2.5x10^5^ MCF7 ESR1^Y357S^ cells were seeded into 10 cm^3^ plates. 48-hour after the initial seeding, each plate was treated with their respective drugs (1 µM Fulv (Selleckchem; Houston, TX, USA; S1191) (dissolved in DMSO)) and/or 1 µM ACSI (EMD Millipore, Burlington, MA, USA; 5.33756) (dissolved in DMSO)) for 24 h. For siRNA experiment, cells were transfected with the siACSS2 and after 24 h of incubation, treated with Veh or Fulv for 24h. We collected and isolated RNA using the same protocol from our previous study ^15^, where the plates were incubated with 1 mL of TRIzol reagent (Life Technologies, Carlsbad, CA, USA) for 15 min on a shaker. After collecting cells from the plate, we mixed them with 250 µL of chloroform and centrifuged. Then, we proceeded to take the clear phase to a new tube and mixed it with 0.5 mL isopropanol. After overnight incubation at 4 °C, we removed the isopropanol and resuspended the RNA in DEPC-treated water. We then used the RNeasy mini kit (Qiagen, Hilden, Germany, cat # 74104) to clean up the RNA. After validation of RNA quantity and purity, samples were submitted to the UIUC sequencing center.

Preprocessing and quality control of data were performed as follows: Fastqc files containing raw RNA-sequencing data were trimmed using Trimmomatic (version 0.38; RRID:SCR_011848) ^24^. Reads were mapped to the human reference genome (GRCh37) from the Ensembl (RRID:SCR_002344) ^25^ database and aligned using the STAR alignment tool (RRID:SCR_015899; version 2.7.0f) ^26^. Read counts were generated from SUBREAD (version 2.0.4) (13), and feature counts were exported for statistical analysis in R. Quality control and normalization were conducted in R using edgeR (version 4.0.12) (14). Statistical analysis was conducted in R using limma (version 3.58.1) (RRID:SCR_010943) (15,16). Empirical Bayesian statistics were conducted on the fitted model of the contrast matrix. Differentially expressed genes were then determined by fold change and P value with a Benjamini–Hochberg multiple testing correction for each gene for each treatment relative to the Veh control. We considered genes with a fold change of >1.5 and P value of <0.05 as significantly differentially expressed.

#### RT-PCR and qPCR

The RNA isolated according to the protocol above was used at 1.5 µg per reaction. For this reaction, we used M-MuLV Reverse Transcriptase (New England Biolabs, M0253). After cDNA synthesis was completed, the cDNA was used to test ESR1 and FASN gene expression. We used GeneCopoeia all-in-one qPCR mix (QP001, GeneCopoeia, Rockville, MD) with ThermoFisher Scientific QuantStudio 7 Pro System (RRID:SCR_020245) from the Biotechnology Center Functional Genomics Facility at UIUC. Primer sequences are provided in the supplementary methods.

### Protein Expression Analysis

#### Western Blot analysis

2.5x10^5^ MCF7 ESR1^Y357S^ cells were seeded into 10 cm^3^ plates on day 0. 24 hours and 48 hours after the initial seeding, each plate was treated with 1 µM Fulv (Selleckchem; Houston, TX, USA; S1191). For the ACSS2 inhibition study, 0.5x10^5^ MCF7 ESR1^Y357S^ cells were seeded into 10 cm^3^ plates. After overnight incubation cells were treated with Veh, 1 µM Fulv, 1 µM ACSI (EMD Millipore, Burlington, MA, USA; 5.33756) (dissolved in DMSO)) or Fulv+ACSI combination for 24 h. For ACSS2 siRNA group, 0.5x10^5^ MCF7 ESR1^Y357S^ cells were seeded into 10 cm^3^ plates. After overnight incubation cells were transfected with ACSS2 siRNA (Santa Cruz Biotechnology, Dallas, TX, USA; sc-72440 and sc-45064) following the manufacturer’s instructions using 50 µM siRNA for 24 h. Then, cells were treated with Veh, 1 µM Fulv for 24h. The cells were collected in lysis buffer, and protein isolation and western blot were performed as described previously ^14^. Briefly, samples were sonicated, and the BCA assay was performed (Thermo Scientific, 23225). Samples were boiled with loading buffer, and 20 µg of protein from each sample was run in 4-20% precast gels (BioRad) and transferred to nitrocellulose membrane. Members were then blocked in blocking buffer (Odyssey, Li-Cor, Lincoln, NE, USA), and ACSS2 and β-actin proteins were probed with AceCS1 antibody ((D19C6) Rabbit mAb #3658 (Cell Signaling Technology, Danvers, MA, USA, RRID:AB_2222710) and β-actin antibody (Sigma-Aldrich, SAB1305546) in 1:1000 and 1:10000 dilutions respectively. The membranes were then visualized with the Licor Odyssey CLx (RRID:SCR_014579) infrared imaging machine and software. The results were normalized for β-actin.

#### Immunofluorescence Assay

5x10^4^ MCF7 ESR1^Y357S^ cells were seeded into 6-well plates on day 0. On Day 2, cell were treated with Veh or 1 µM Fulv for 24 hours. Next, cells were fixed in 4% paraformaldehyde for 15 minutes. 10 minute 0.5% Triton X-100 incubation was used to permeabilize the cell membrane. For ACSS2 detection, AceCS1 antibody (D19C6) Rabbit mAb #3658 (Cell Signaling Technology, Danvers, MA, USA; RRID:AB_2222710) was used at 1:100 dilution. As secondary antibody, Goat anti-rabbit IgG (H+L) Highly Cross-Adsorbed Secondary Antibody, Alexa Fluor™ 488 (Thermo Fisher Scientific Cat# A-11034, RRID:AB_2576217) was used at 5 µg/mL concentration. Before visualization, SlowFade™ Gold antifade reagent with DAPI (Thermo Fisher Scientific, Waltham, MA, USA) was applied to visualize the nucleus. The EVOS M500 Imaging System (Thermo Fisher Scientific, Waltham, MA, USA, RRID:SCR_023650) instructions were followed for visualization and quantitative analysis.

### Cell Viability Assays

#### Acetyl-CoA Acetyltransferase 1 (ACAT1) Inhibition

1 × 10^3^ MCF7 ESR1^Y537S^ cells were seeded in each well of a 96-well plate on day 0. On day 1, cells were treated with 1 µM ATR-101 (ACAT1 inhibitor; Sigma-Aldrich, St. Louis, MO, USA) with and without 1 µM Fulv. On day 4, treatment was repeated. On Day 7, the WST-1 Assay (Millipore Sigma, Burlington, MA, USA) was performed. Cell viability was measured using a Cytation 5 cell imaging multimode reader (RRID:SCR_019732) at an optical density of 450 nm.

#### Acetate Dose Response

1x10^3^ MCF7 ESR1^Y357S^ cells were seeded into 96-well plates on day 0. On day 1, cells were treated with 0, 50, or 100 µM of acetate (Sigma Aldrich; Burlington, MA, USA; PHR2615-2G) (dissolved in water) in the presence or absence of 1 µM Fulv (n=12). On day 4, treatment was repeated. On Day 7, the WST-1 Assay (Millipore Sigma, Burlington, MA, USA) was performed. Cell viability was measured using a Cytation 5 cell imaging multimode reader (RRID:SCR_019732) at an optical density of 450 nm.

#### Fatty Acid Loading

1x10^3^ MCF7 ESR1^Y357S^ cells were seeded into 96-well plates on day 0. On day 1, cells were treated with 0, 0.01, 0.1, or 1 µM of palmitate (Cayman Chemical; Ann Arbor, Michigan; 10006627) (dissolved in ethanol) in the presence or absence of 1 µM Fulv (n=12). On day 4, treatment was repeated. On Day 7, the WST-1 Assay (Millipore Sigma, Burlington, MA, USA) was performed. Cell viability was measured using a Cytation 5 cell imaging multimode reader (RRID:SCR_019732) at an optical density of 450 nm.

#### ACSS2 siRNA Transfection

The experiment comprised eight groups with six replicates each: Veh, Fulv, ACSS2 inhibitor (ACSI), Fulv+ACSI, siRNA, siRNA+Fulv, siRNA+ACSI, and siRNA+Fulv+ACSI (1 µM Fulv and/or 1 µM ACSI). 1x10^4^ MCF7 ESR1^Y357S^ cells were seeded into 96 well plates in DMEM media with 10% HI-FBS. Next day, the cells were transfected with ACSS2 siRNA (Santa Cruz Biotechnology, Dallas, TX, USA; sc-72440 and sc-45064) following the manufacturer’s instructions using 50 µM siRNA. After overnight incubation, cells were treated with Veh or 1 µM Fulv. Treatment was repeated 48 h later. On day 7, cell viability assay using WST-1 was performed. Optical density at 450 nm with Cytation 5 cell imaging multimode reader (RRID:SCR_019732) was measured.

### Isotope Tracing Assay

The experiment consisted of 12 groups with 3 replicates each, divided into two main categories:

i 13C-labeled acetate groups (Cambridge Isotope Laboratories, Inc., Tewksbury, MA, USA; CLM-440-PK): Veh, Fulv, ACSI, Fulv+ACSI, ACSS2 siRNA, ACSS2 siRNA+Fulv
ii 12C-unlabeled acetate groups (Sigma Aldrich, Burlington, MA, USA; PHR2615-2G): Veh, Fulv, ACSS2 inhibitor (ACSI), Fulv+ACSI, siRNA, siRNA+Fulv

2.5x10^5^ MCF7 ESR1^Y357S^ cells were seeded to 10 cm^3^ plates with DMEM media with 10% HI-dialyzed FBS (Gibco, USA; A3382001). The siRNA groups were transfected with ACSS2 siRNA the next day following the manufacturer’s protocol. The following day, Fulv, ACSI, or Fulv+ACSI treatments were given for 24h (1 µM Fulv and/or 1 µM ACSI). After 24h incubation with drugd, 100 µM labeled/unlabeled acetate was fed to cells. After 30 minutes of acetate exposure, we collected the cells and submitted them to the UIUC Metabolomics Center for measurement of labeled carbon levels using liquid chromatography-mass spectrometry (LC-MS).

### Epigenomic Analysis

#### CUT&RUN Sequencing

Cleavage under targets and release using nuclease (CUT&RUN) is an epigenomic profiling method that separates specific protein-bound DNA ^27^. For this assay, 2.5 × 10^5^ MCF7 ESR1^Y537S^ cells were seeded in media described above to 10 cm^3^ plates on day 0. Four groups were prepared: Vehicle, Fulv, ACSI, and Fulv+ACSI, with 3 replicates in each group. Cells were treated with 1 μM ACSI for 24h on day 2 and followed by 45 minutes of 1μM Fulv treatment on day 3. After both treatments, cells were collected in cell culture media using trypsin. CUT&RUN assay was performed using the CUT&RUN assay kit (Cell Signaling Technologies, #86652, MA, USA) according to the manufacturer’s instructions. Chromatin enrichment of three factors was tested: ER⍰ and ACSS2, along with one histone acetylation: H3K27ac.

For each sample, 1 × 10^5^ MCF7 ESR1^Y537S^ cells were used. Proteins were crosslinked with 0.1% formaldehyde for 2 minutes at room temperature (RT). Crosslinking was stopped by 1X glycine solution treatment for 5 minutes at RT. After washing, the cells were treated with concanavalin A beads and subsequently with their respective antibodies at 4°C overnight. An ACSS2 antibody cocktail (2 µL each of Cell Signaling Technology Cat# 3658, RRID:AB_2222710 and Thermo Fisher Scientific Cat# MA5-14810, RRID:AB_10983922), H3K27ac antibody (2 µL, Abcam Cat# ab4729, RRID:AB_2118291), and ER⍰ antibody cocktail (30 µL each of Santa Cruz Biotechnology Cat# sc-8002, RRID:AB_627558 and Santa Cruz Biotechnology Cat# sc-8005, RRID:AB_627556) were used. The cells were then treated with digitonin and pAG-MNase enzyme. The enzyme was activated with cold calcium chloride, and DNA fragments were released by stop buffer incubation. Finally, the enriched chromatin samples were treated with SDS and proteinase K to reverse the crosslinking. After DNA purification using a kit specifically designed for CUT&RUN (Cell Signaling Technologies, #14209, MA, USA), the samples were sent to the UIUC sequencing center for further analysis.

#### ATAC Sequencing

Similar to CUT&RUN sequencing, we prepared 4 groups: Vehicle, Fulv, ACSI, and Fulv+ACSI, and had 3 replicates in each group. Cells were treated with ACSI for 24 hours and Fulv treatment for 45 min. Cells were collected, and 1x10^5^ live cells were submitted to the UIUC sequencing center.

### Preprocessing and Quality Control

#### CUT&RUN Sequencing

48 samples were submitted to High-Throughput Sequencing and Genotyping Unit in the Roy J Carver Biotechnology Center (CBC) at UIUC to generate CUT&RUN libraries. We used Bowtie2 (v2.4.5) (RRID:SCR_016368) to align paired-end FASTQ files to human reference (GRCh38). Mapped reads were further sorted and indexed using SAMtools (v1.17) (RRID:SCR_002105) to prepare individual as well as merged BAM files. Narrow and broad peaks were called for these BAM files using MACS2 (v2.2.5) (RRID:SCR_013291) with input reads as control. For ACSS2 and ER⍰ samples, we used narrow peaks, and for H3K27ac, we used broad peaks. Depending on the analysis type, we used merged or individual files.

#### ATAC Sequencing

12 samples were submitted to High-Throughput Sequencing and Genotyping Unit in the Roy J Carver Biotechnology Center (CBC) at UIUC to generate ATAC-Seq libraries. Read alignment and peak calling were performed using nf-core/atacseq (version 2.1.2) workflow. Briefly, reads were aligned to GRCh38 human genome assembly using the BWA aligner (RRID:SCR_010910), and broad peaks were called using MACS2 software (RRID:SCR_013291).

### Statistical Analysis

Data from the retrospective study was tested for normal distribution, and outliers were removed using the Robust regression and Outlier removal (ROUT) method. For normally distributed data, we used unpaired Welch’s t-tests and one-way ANOVA to identify significant differences between groups (*p < 0.05; **p < 0.01; ***p < 0.001; ****p < 0.0001). We used GraphPad Prism version 9 for Mac (GraphPad Software, https://www.graphpad.com GraphPad Prism (RRID:SCR_002798)) to prepare survival plots.

Data from cell viability assays were analyzed using Microsoft Office Excel for Mac and GraphPad Prism version 9 for Mac (GraphPad Software, https://www.graphpad.com GraphPad Prism (RRID:SCR_002798)). For the isotope tracing assay, Non-Targeted Tracer Fate Detection (NTFD) software was employed, and visualization was performed using GraphPad Prism. For CUT&RUN and ATAC sequencing analyses, heatmaps were prepared using deepTools computeMatrix and plotHeatmap with Galaxy tools (https://usegalaxy.org/, RRID:SCR_006281) with merged bam files. Gene annotation was performed using GREAT version 4.0.4 (http://great.stanford.edu/public/html/, RRID:SCR_005807). Binding site intensities were visualized with the UCSC Genome Browser (https://genome.ucsc.edu/, RRID:SCR_005780), and binding site distribution was determined using ChIPseeker (RRID:SCR_021322) from Galaxy. The top 1% and 10% binding magnitude changes for ER⍰ and ACSS2 bindings respectively, were determined using the peaks data.

Data from the immunofluorescence assays were analyzed using Fiji (RRID:SCR_002285) ^28^ to determine the difference in the nuclear and cytosolic ACSS2 levels and visualized using GraphPad Prism (RRID:SCR_002798).

Homer (RRID:SCR_010881) was applied for motif analysis, and Gene Set Enrichment Analysis (GSEA, RRID:SCR_003199, https://www.gsea-msigdb.org/) was used for gene set determination. Visualization of these results was carried out using GraphPad Prism (RRID:SCR_002798).

For RNA sequencing, the same technique as in the previous study ^15^ was utilized. Briefly, Cluster3 software was used to cluster differentially expressed genes. k-means clustering with Euclidean distance similarity metric is used to organize genes into 40 clusters with 100 runs and data were visualized using Java Treeview (RRID:SCR_016916). Gene set enrichment analysis (RRID:SCR_003199) ^29,30^ was used to identify Gene Ontology terms associated with different treatments. Correlation analysis was also performed for this data using GraphPad Prism (RRID:SCR_002798). Ridgeline plots of enriched metabolic pathways based on RNA sequencing results comparing different groups in various metabolic pathways are prepared with ExpressAnalyst (https://www.expressanalyst.ca). For all data and the qPCR results, one-way and two-way ANOVA tests were applied to compare different condition effects, and these analyses were conducted using GraphPad Prism (RRID:SCR_002798).

In vivo study results were analyzed using the signal intensity obtained from weekly IVIS imaging to measure the progression of the metastatic tumors. The outliers were determined and removed using the GraphPad ROUT method. Each mouse’s tumor signal intensity at the beginning of the treatment (Day 32) was assigned as 100%, and the increase in metastatic burden on the last day of treatment (Day 56) was calculated in proportion. The change was analyzed using unpaired t-test of each treatment relative to Veh.

## 4. Results

### Liver Metastasis is Associated with Poor Prognosis and Reduced Fulv Efficacy in Breast Cancer Patients

Our cohort had 192 patients that satisfied the inclusion criteria. Of these 192 patients, 60 (31.2%) had liver metastasis while 132 (68.8%) had metastasis to another site. Only five patients (2.6%) were male (**Table 1**). Analysis of metastatic patterns revealed that bone metastasis was the most common (69.8%), followed by liver metastasis (31.3%) and lymph node metastasis (31.3%) (**Fig. 1A-left**). Only 13.3% of liver metastatic patients had single-site metastasis, while 86.7% of liver metastatic patients had other accompanying metastasis, most commonly bone and lymph nodes (**Fig. 1A-right, 1B**). Patients with liver metastases demonstrated significantly worse clinical outcomes compared to those with non-liver metastases. The mortality rate was substantially higher in the liver metastasis group (83.3%) versus the non-liver metastasis group (62.7%) (**Fig. 1C**). Survival analysis confirmed significantly reduced survival in liver metastatic patients both from initial breast cancer diagnosis (HR 2.653, 95% CI 1.708-4.123, p=0.0006) and from metastasis diagnosis (HR 1.997, 95% CI 1.304-3.058, p=0.0043) (**Fig. 1D**). Additionally, patients with liver metastases experienced a significantly shorter disease-free interval, with faster progression from initial breast cancer diagnosis to first metastatic presentation compared to patients with other metastatic sites (p=0.0312) (**Fig. 1E**). Analysis of comorbidities revealed no statistically significant differences between liver metastatic and non-liver metastatic groups for underlying conditions including lipoprotein metabolism disorders, hypertension, diabetes, chronic kidney disease, and ischemic heart disease, either before or after metastasis diagnosis (**Suppl. Fig. 1, Table 1**). However, Fulv treatment efficacy showed striking disparities between groups. Among Fulv-treated patients, survival rates were markedly lower in the liver metastasis group (7.4%) compared to the non-liver metastasis group (43.9%) (**Fig. 1F**), indicating that Fulv treatment is significantly less effective in improving overall survival for patients with liver metastases compared to those with metastases at other sites.

**Table1.** Patient characteristics.

**Figure 6**
|  | **All** | **Liver Metastatic** | **Other Metastatic** |
| --- | --- | --- | --- |
| Number of patients | 192 | 60 | 132 |
| Female patients | 187 | 60 | 127 |
| Male patients | 5 | 0 | 5 |
| Lipidemia before mets Dx | 105 | 30 | 75 |
| Lipidemia after mets Dx | 11 | 4 | 7 |
| HT before mets Dx | 98 | 22 | 76 |
| HT after mets Dx | 14 | 4 | 10 |
| DM before mets Dx | 37 | 8 | 29 |
| DM after mets Dx | 21 | 6 | 15 |
| CKD before mets Dx | 10 | 5 | 5 |
| CKD after mets Dx | 14 | 5 | 9 |
| IHD before mets Dx | 16 | 3 | 13 |
| IHD after mets Dx | 19 | 4 | 15 |

**Figure 1.**
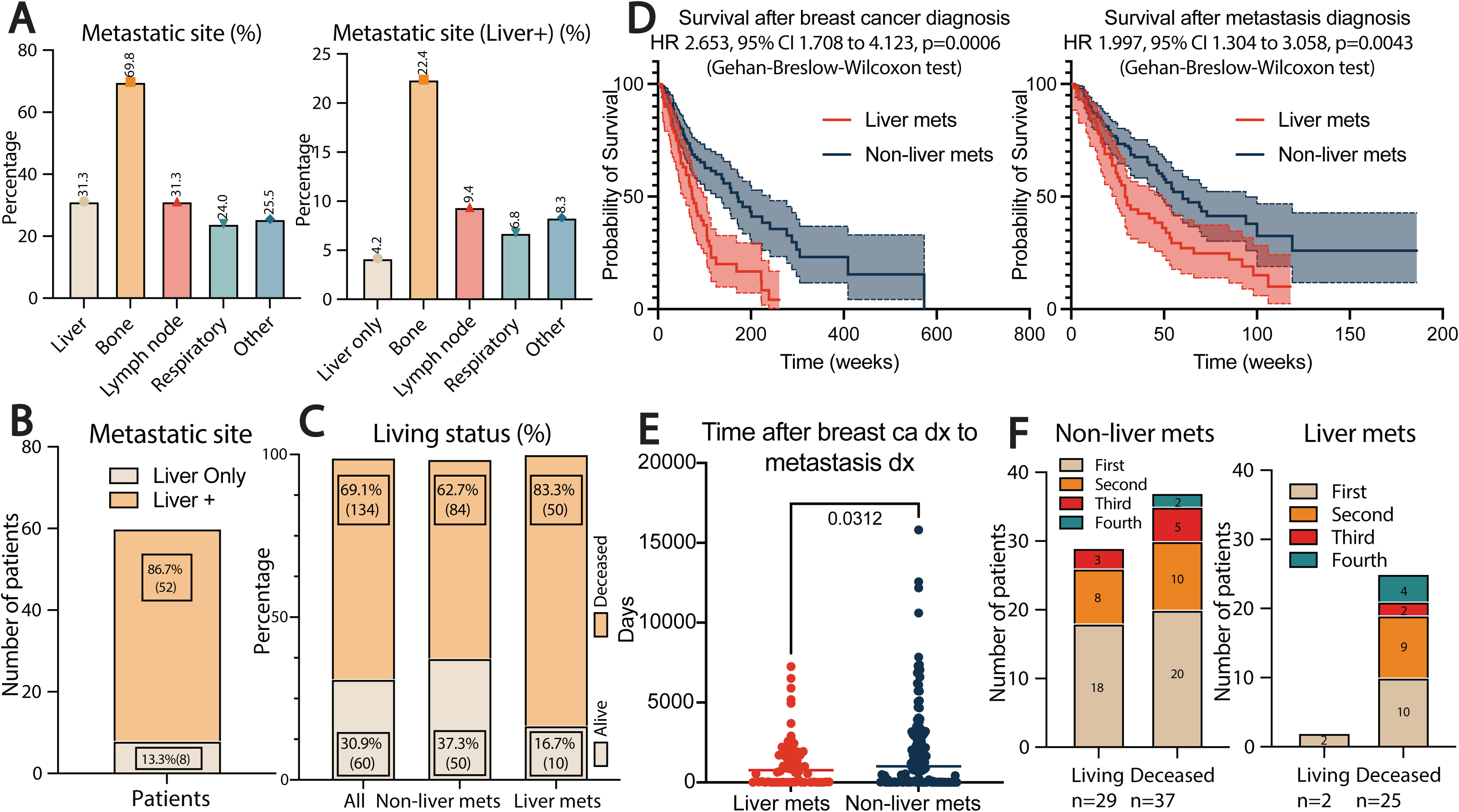
Liver Metastasis is Associated with Poor Prognosis and Reduced Fulv Efficacy in Breast Cancer Patients. **(A)** Distribution of metastatic sites in the study cohort, shown as percentages (left) and distribution of concurrent metastatic sites in patients with liver metastases, shown as percentages (right). **(B)** Distribution of patients with liver-only metastases versus liver metastases with additional sites. **(C)** Survival status stratified by metastatic site (all patients, non-liver metastasis group, and liver metastasis group). **(D)** Kaplan-Meier survival curves comparing liver and non-liver metastasis groups after breast cancer diagnosis (top) and metastasis diagnosis (bottom). **(E)** Time interval between breast cancer diagnosis and metastasis diagnosis in liver versus non-liver metastasis groups. **(F)** Treatment line distribution by survival status for patients with non-liver metastases (left) and liver metastases (right) receiving Fulv therapy.

### Fulv Treatment Upregulates Fatty Acid Metabolism and Cancer Progression Pathways in Liver Metastasis-Derived PDX Model

To examine molecular changes induced by Fulv in a Fulv resistant model, we used HCI044EI PDX, which was previously reported to metastasize to liver ^23^. While growth of a commonly used, endocrine therapy responsive PDX, HCI013 was completely blocked by Fulv treatment (**Supp. Fig. 1B**), HCI044EI PDX tumors continued to grow in the presence of Fulv treatment without significant changes to animal weights (**Fig. 2A and 2B**). Thus, this model enabled us to focus on metabolic and gene expression changes induced by Fulv in an endocrine therapy resistant model. Metabolomics analysis of HCI044EI PDX samples examining the changes induced by Fulv revealed significant alterations in the metabolic profile following Fulv treatment, with changes particularly enriched in fatty acid metabolism, ketone body metabolism, and phospholipid biosynthesis pathways (**Fig. 2C, 2D**). The metabolite heatmap demonstrated distinct clustering of metabolites that change between Veh-and Fulv-treated tumors, indicating substantial metabolic reprogramming in response to endocrine therapy (**Fig. 2C**). To further characterize these molecular changes, we performed targeted gene expression analysis using the NanoString Human Metabolic panel on PDX tumor samples. Heatmap visualization of significantly altered genes revealed distinct expression clusters between treatment groups (**Fig. 2E**). Fulv treatment upregulated multiple pathways, including nuclear receptor signaling, chromatin modification, and cancer-related processes (**Suppl. Table 1**). Pathway analysis demonstrated that Fulv treatment enhanced expression of genes associated with key cancer hallmarks, particularly reprogramming of energy metabolism, replicative immortality, resistance to cell death, genomic instability, sustained angiogenesis, and tissue invasion and metastasis (**Fig. 2E, right panel**). We further validated Fulv regulation of select genes using quantitative RT-PCR, including Ki67, FOXM1, HADHB, and G6PD (**Fig. 2F**).

**Figure 2.**
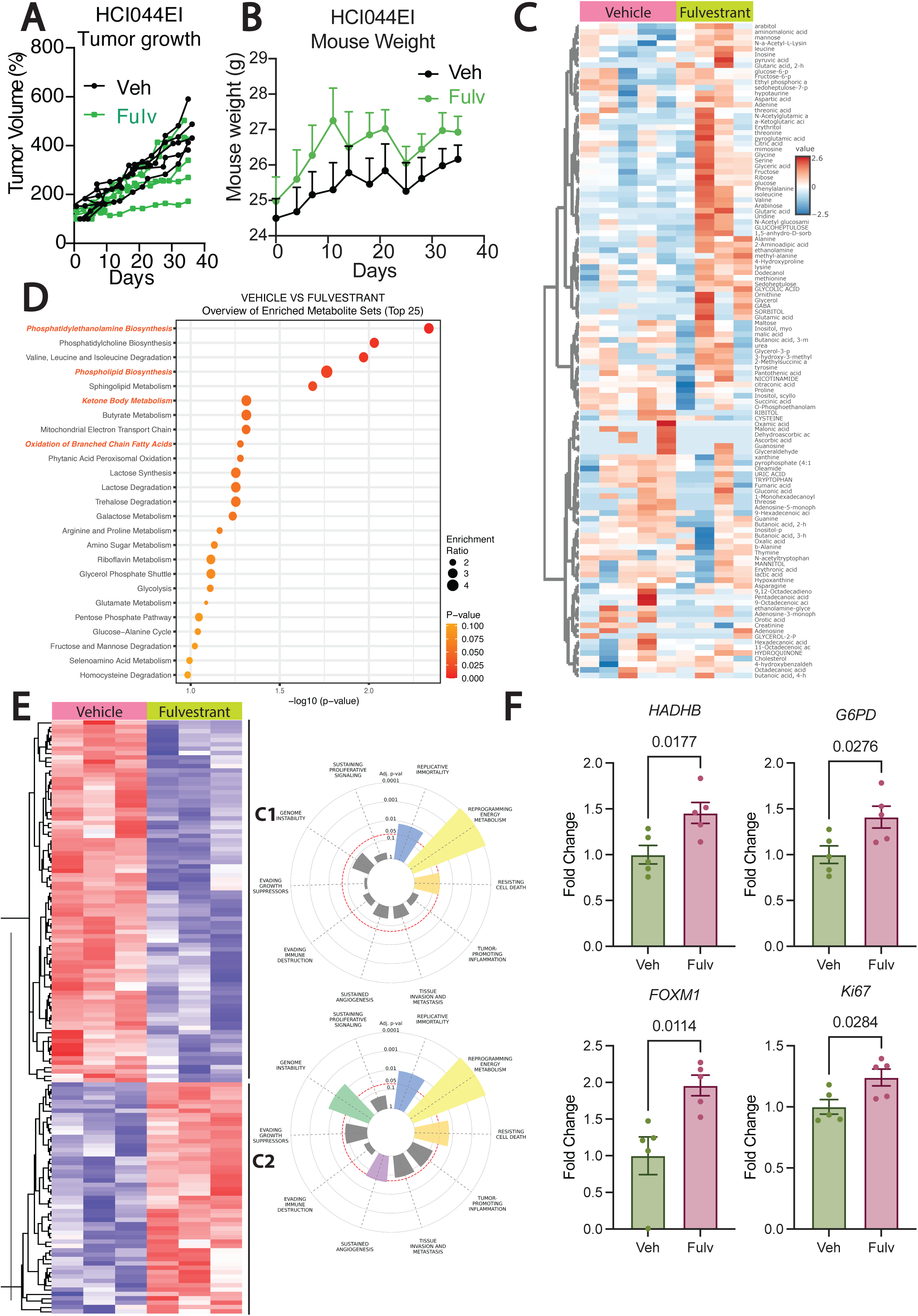
Fulv Treatment Upregulates Fatty Acid Metabolism and Cancer Progression Pathways in liver metastasis-derived PDX model. **(A)** The changes in tumor volume and **(B)** the changes in mouse weight with and without Fulv treatment for HCI044EI (N=5 per group) PDX tumors. **(C)** Heatmap for the metabolomics assay showing non-treated (Veh) (N=5) and Fulv-treated (N=4) HCI044EI PDX tumors. **(D)** Overview of enriched metabolite sets showing enriched metabolites in Fulv-treated HCI044EI PDX tumors compared to Veh-treated. **(E)** Heatmap for the Nanostring assay showing only the significantly altered gene expression profile based on multiple t-test for HCI044EI PDX tumors comparing Veh and Fulv groups **(left)**, cancer hallmarks enrichment for the genes in each cluster in the heatmap **(right)** (N=3 per group). **(F)** qPCR for various genes with statistically significant changes (n=5 per group).

### Fulv Treatment Upregulates ACSS2 Expression and Acetate-Derived Acetyl-CoA Synthesis in Metastatic Breast Cancer Cells

Based on our previous findings that *ACSS2* was upregulated in Fulv-treated cell line xenograft tumors^15^, a small molecule inhibitor of ACSS2, ACSI synergized with Fulv to reduce cell viability in MBC cells ^15^ and the observed upregulation of fatty acid metabolites in HCI044EI PDX model following Fulv treatment (**Fig. 2C**), we focused on ACSS2 due to its critical roles in fatty acid synthesis, ketogenesis, and transcriptional regulation (**Fig. 3A**). Consistent with our in vivo results from xenografts ^15^, we observed a similar trend toward increased *ACSS2* mRNA expression in MCF7-ESR1^Y537S^ cells treated with Fulv in vitro (**Fig. 3B**). The alteration we observed in mRNA expression is more pronounced and statistically significant at the protein level, as protein expression of ACSS2 continued to increase throughout 48 hours of Fulv treatment (**Fig. 3C**). While siRNA targeting of ACSS2 resulted in more than 60% reduction in protein level, treatment with ACSI did not affect ACSS2 protein levels (**Fig. 3D**). This increase in ACSS2 expression was accompanied by enhanced cell viability in the presence of Fulv and increasing dose of acetate (**Fig. 3E**) and metabolic activity, as demonstrated by elevated acetyl-CoA production measured by isotope tracing with ^13^C-labeled acetate (**Fig. 3F**). While siRNA targeting of ACSS2 significantly reduced acetyl-CoA production upon Fulv treatment, ACSI treatment reduced only the basal level, potentially due to shorter inhibition of ACSS2 during this treatment. These findings support that Fulv treatment promotes acetyl-CoA synthesis through the ACSS2-mediated acetate utilization pathway. Given the observed metabolic reprogramming, we investigated whether targeting ACSS2 could restore Fulv sensitivity in MBC cells. We targeted ACSS2 protein and activity and examined the impact on Fulv-induced cell viability over a week. Both metabolic inhibitions using ACSS2 inhibitor (ACSI) and genetic silencing through ACSS2 siRNA significantly enhanced Fulv efficacy, reducing cell viability compared to Fulv treatment in control cells (**Fig. 3G**). These results demonstrate that Fulv treatment induces a metabolic adaptation characterized by increased ACSS2 expression and enhanced acetate-derived acetyl-CoA synthesis. Targeting this adaptive response through ACSS2 inhibition represents a promising strategy to overcome endocrine resistance and improve therapeutic efficacy in metastatic breast cancer.

**Figure 3.**
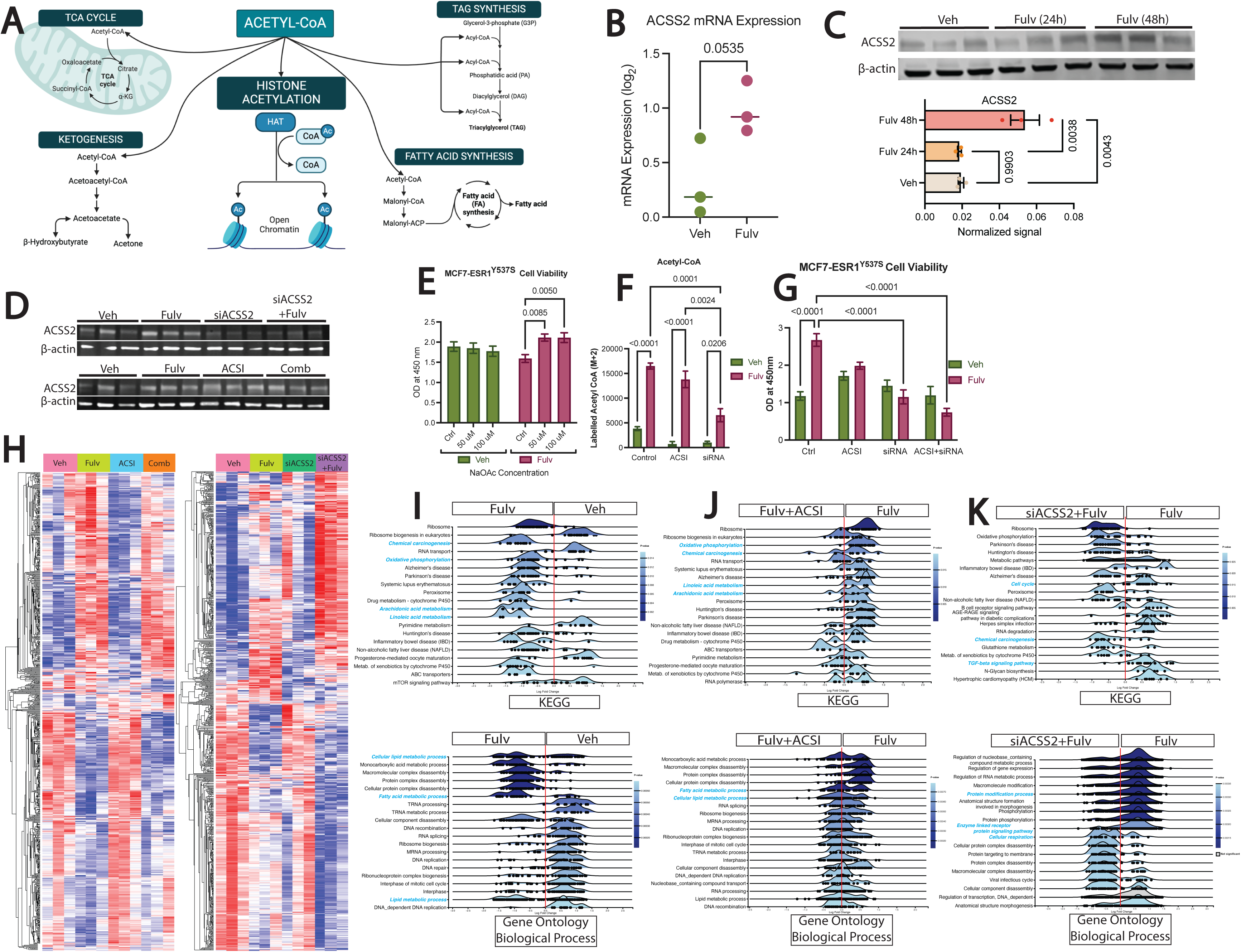
Fulv Treatment Upregulates ACSS2 Expression and Acetate-Derived Acetyl-CoA Synthesis in Metastatic Breast Cancer Cells. **(A)** The metabolic pathways in which acetyl-CoA is involved. **(B)** Changes in the ACSS2 expression with Veh and Fulv treatment groups in vitro (n=3 per group). **(C)** Western blot bands for ACSS2 and β-actin with no treatment (Veh), 24-hour, and 48-hour Fulv treatments (top) and bar graphs displaying the differences between these groups for normalized signals (bottom). **(D)** Western blot analysis for ACSS2 and β-actin with no treatment (Veh), Fulv, ACSS2 siRNA treatment and Fulv + ACSS2 siRNA treatment (top) and no treatment (Veh), Fulv, ACSS2 inhibitor (ACSI) treatment, and combination of Fulv with ACSS2 inhibitor (ACSI) treatment (bottom). **(E)** Cell viability findings for increasing acetate levels (0-50-100 µM) with Fulv treatment (n=16 per group). **(F)** Changes in the acetyl-CoA levels with and without pharmacological and genetic ACSS2 inhibition and Fulv treatment (n=3 per group). **(G)** Cell viability findings with metabolic and/or genetic ACSS2 inhibition with Fulv treatment (n=6 per group). **(H)** Heatmap of gene expression profiles for Veh, Fulv, ACSI, and Fulv+ACSI treatments (left) and for Veh, Fulv, ACSS2 siRNA, and Fulv+ACSS2 siRNA treatments (right) (n=3 per group). Ridgeline plot of enriched metabolic pathways based on RNA sequencing results comparing **(I)** Veh and Fulv, **(J)** Fulv and Fulv+ACSI Comb, and **(K)** Fulv and Fulv+siACSS2 Comb in KEGG (left) and Gene Ontology: Biological Process (right) pathways. These graphs were prepared with ExpressAnalyst (https://www.expressanalyst.ca).

To examine the functional consequences inhibiting ACSS2, we performed genome-wide mRNA profiling to assess how ACSS2 depletion (siRNA) or inhibition (ACSI) affects Fulv-mediated gene expression changes. RNA sequencing analysis revealed differences between treatment groups, with both ACSI (**Fig. 3H-left**) and ACSS2 siRNA (**Fig. 3H-right**) substantially altering the Fulv-induced transcriptional landscape. Pathway analysis of genes upregulated by Fulv treatment compared to Veh revealed significant enrichment in oxidative phosphorylation, arachidonic acid and linoleic acid metabolism pathways, as well as cellular lipid metabolic, fatty acid metabolic, and general lipid metabolic processes (**Fig. 3I**). These findings confirm that Fulv treatment promotes metabolic reprogramming toward lipid synthesis and energy metabolism, consistent with our PDX results. Critically, comparison of Fulv+ACSI combination therapy to Fulv monotherapy demonstrated that ACSS2 inhibition blocked the Fulv-mediated upregulation of these metabolic pathways, including oxidative phosphorylation, arachidonic acid and linoleic acid metabolism, and lipid metabolic processes (**Fig. 3J**). Additionally, ACSS2 inhibition prevented Fulv-induced activation of cell cycle and TGF-β signaling pathways, as well as protein modification and enzyme-linked receptor signaling pathways (**Fig. 3K**). These results demonstrate that Fulv stimulates fatty acid synthesis and cell proliferation-related gene expression in an ACSS2-dependent manner, and that targeting ACSS2 can reverse this transcriptional reprogramming.

### ACSS2-Mediated Acetate Incorporation Drives Metabolic Reprogramming Following Fulv Treatment

To validate functional consequences of transcriptional changes we observed with ACSS2 inhibition, we treated MCF7-ESR1^Y537S^ cells with Fulv and/or ACSI, which resulted in significant alterations in the cellular metabolic profile (**Fig. 4A**). Metabolic pathway analysis revealed that Fulv treatment significantly upregulated pathways involved in cell proliferation, including the pentose phosphate pathway and purine and pyrimidine metabolism. Additionally, Fulv enhanced lipid metabolism-related pathways, including CoA biosynthesis, linolenic and linoleic acid metabolism, and steroid biosynthesis compared to Veh treatment (**Fig. 4B-top**). Comparison of Fulv monotherapy to combination treatment confirmed that lipid metabolism pathways (steroid biosynthesis, CoA metabolism) and cell proliferation pathways (purine/pyrimidine metabolism, pentose phosphate pathway) were significantly upregulated with Fulv alone (**Fig. 4B-bottom**). To directly assess acetate incorporation into metabolic pathways via ACSS2, we performed stable isotope tracing using ^13^C-labeled acetate in cells with pharmacological or genetic ACSS2 inhibition. ACSS2 expression and activity significantly influenced the levels of acetate-labeled metabolites across all treatment conditions. Detailed metabolic flux analysis revealed that Fulv treatment redirected acetate flux toward lipid synthesis, (**Fig. 4C**) and away from the TCA cycle as evidenced by decreased carbon incorporation into TCA cycle intermediates like oxalic acid (**Fig. 4D**). Of note, none of these metabolites in **Fig. 4C** had ^13^C incorporation when ACSS2 activity was depleted using siRNA or ACSI. These findings align with our PDX model results, where the Fulv-resistant HCI044EI line showed upregulated fatty acid synthesis pathways. While our previous work showed increased *ACAT1* expression, which converts acetyl-CoA to acetoacetyl-CoA, with Fulv treatment ^15^, ACAT1 inhibition did not reduce cell viability (**Fig. 4E**). Furthermore, palmitate supplementation to bypass potential fatty acid synthesis deficits did not rescue the reduced cell viability observed with ACSI + Fulv combination treatment or ACSS2 kock-down (**Fig. 4F**), indicating that fatty acid synthesis is not the primary pathway mediating ACSS2-associated Fulv resistance. These findings demonstrate that while Fulv treatment promotes acetate flux toward lipid synthesis, the therapeutic benefit of ACSS2 inhibition is not primarily due to impaired fatty acid metabolism. This led us to investigate another critical acetyl-CoA-dependent process: epigenetic regulation through histone acetylation.

**Figure 4.**
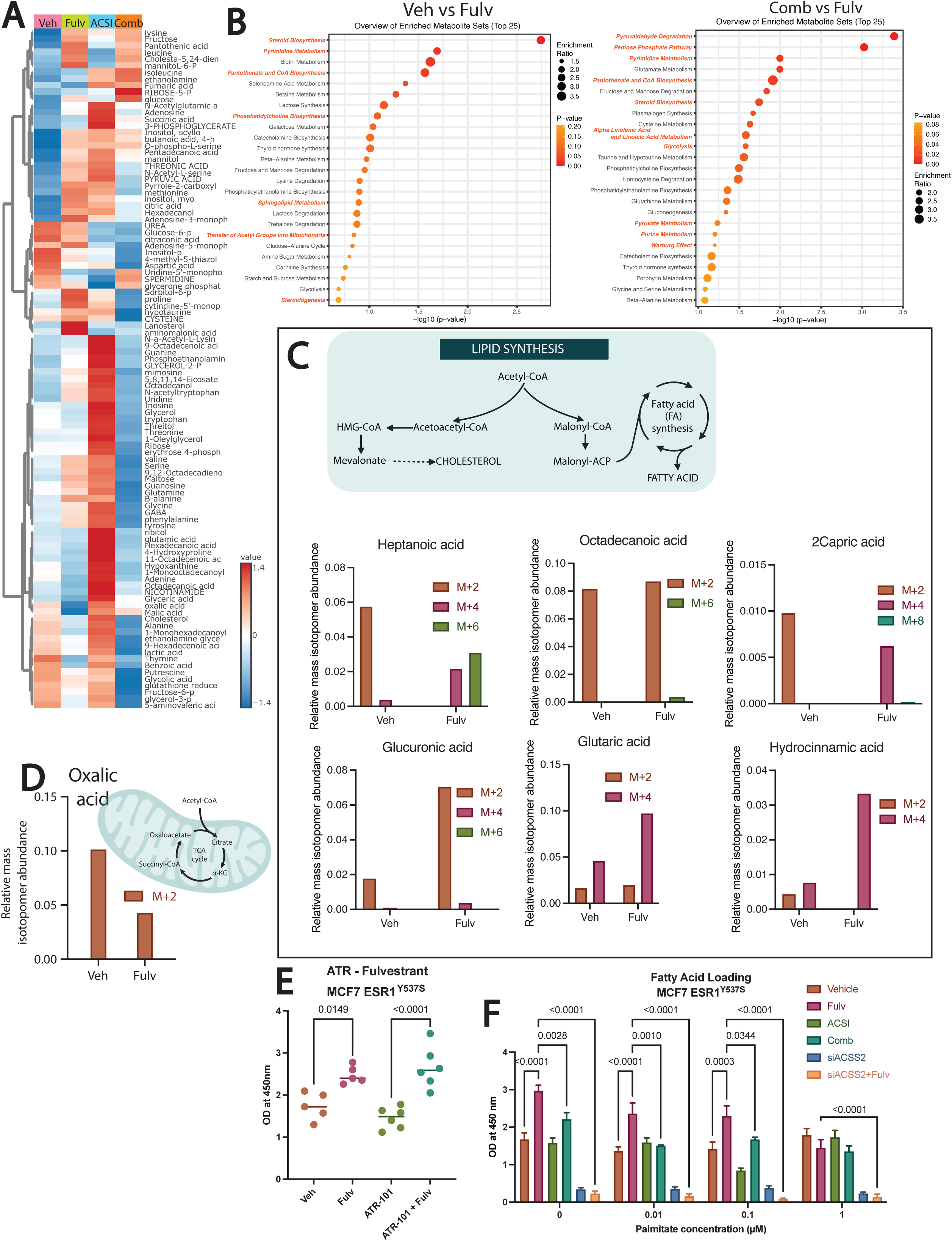
ACSS2-Mediated Acetate Incorporation Drives Metabolic Reprogramming Following Fulv Treatment. **(A)** Heatmap showing metabolomics findings with Fulv and/or ACSI treatments (n=3 per group). **(B)** Overview of enriched metabolite sets showing enrichment for Fulv compared to Veh (top) and enrichment for Fulv compared to the combination treatment (bottom). **(C)** Levels of labeled carbons from metabolites in fatty acid synthesis with Veh and Fulv treatment. **(D)** Levels of labeled carbons from metabolites in the TCA cycle with Veh and Fulv treatment. **(E)** Cell viability assay results with Fulv and ATR-101 (ACAT1 inhibitor) ^15^. **(F)** Cell viability levels with increasing palmitate concentrations (0, 0.01, 0.1, 1 µM) with different treatments (Fulv, ACSI, Fulv+ACSI Comb, siACSS2, siACSS2+Fulv).

### ACSS2 Nuclear Localization and Chromatin Co-occupancy with ER**α** Drive Fulv Resistance

To examine the epigenetic role of ACSS2, we first characterized its cellular localization using immunofluorescence analysis. This analysis revealed that ACSS2 was mainly nuclear in MCF7 parental cells as well as cells with ESR1 mutations (**Fig. 5A**), and nuclear signal for ACSS2 further increased in MCF7-ESR1^Y537S^ mutant cells following Fulv treatment, whereas no significant changes were observed in non-mutated MCF7 parental cells or MCF7-ESR1^D538G^ mutant cells (**Fig. 5A**). Importantly, ACSS2 showed predominantly nuclear localization in both MCF7-ESR1^D538G^ and MCF7-ESR1^Y537S^ cells, contrasting with the more cytoplasmic distribution observed in MCF7 parental cells (**Fig. 5B**). This nuclear enrichment supports a potential epigenetic regulatory role for ACSS2 through local acetyl-CoA provision for histone acetyltransferases. To examine a potential role for ACSS2 in ERα-mediated gene expression, we performed CUT&RUN analysis for ACSS2, ERα and H3K27ac histone mark. CUT&RUN analysis revealed distinct chromatin binding patterns for ACSS2, ERα, and H3K27ac under different treatment conditions (Vehicle, Fulv, ACSI, Fulv+ACSI) (**Fig. 5C**). The most striking finding was increased ACSS2 chromatin recruitment following Fulv treatment, which was completely abolished by ACSI treatment. ATAC-seq analysis showed no changes in global DNA accessibility for these factors (**Suppl. Fig. 2**), indicating that ACSS2 activity exerts localized rather than genome-wide effects on DNA accessibility. However, H3K27 acetylation patterns changed significantly across treatments, consistent with local modulation of histone acetylation rather than global changes. Critically, the number of overlapping binding sites for ACSS2, ERα, and H3K27ac increased dramatically with Fulv treatment but decreased substantially with combination therapy (**Fig. 5D**, **Table 2**). Since combination treatment also reduces cell viability, these co-occupied sites likely play essential roles in cell survival and proliferation. Analysis of binding intensity revealed that both ERα (top 1%) and ACSS2 (top 10%) binding increased with Fulv treatment and decreased with combination therapy, with ACSS2 showing more pronounced changes (**Fig. 5E**), demonstrating the significant impact of these treatments on chromatin occupancy.

**Figure 5.**
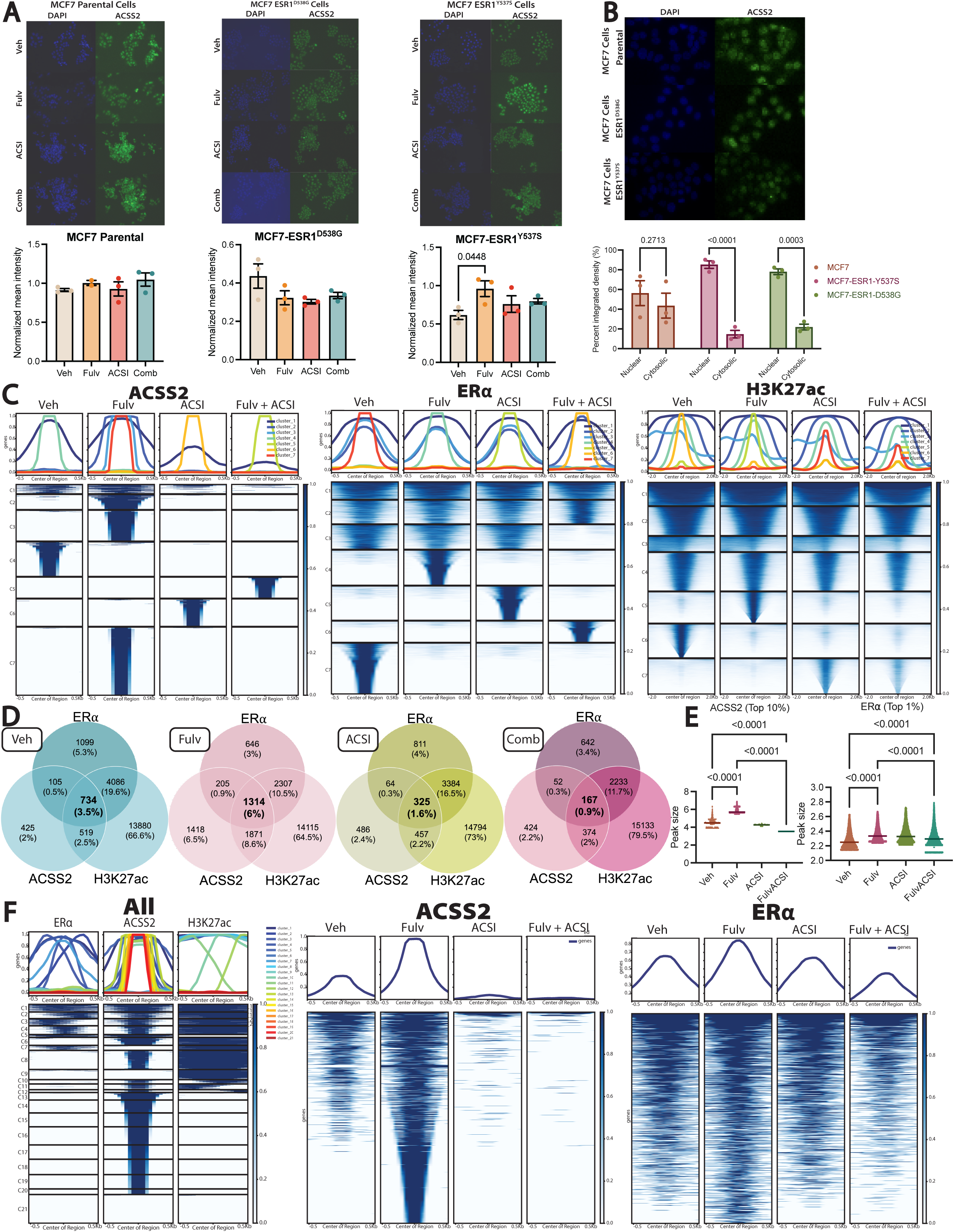
Nuclear ACSS2 Co-localizes with ERα and H3K27ac to Drive Fulv-Induced Epigenetic Reprogramming. **(A)** Immunofluorescence images and signal intensity plots showing ACSS2 levels for MCF7 parental (non-mutated) cells (left), MCF7 ESR1^D538G^ cells (middle), and MCF7 ESR1^Y537S^ cells (right) (n=3 per group). **(B)** Immunofluorescence images and the integrated density plot obtained from Fiji software for the intracellular localization of ACSS2 protein for MCF7 ESR1^Y537S^ cells (n=3 per group). **(C)** Heatmaps for ACSS2 binding (left), ER⍰ binding (middle) and H3K27 acetylation (right) pulldowns (n=3 per group). **(D)** Venn diagram showing the number of binding sites exclusive to each treatment (Veh, Fulv, ACSI, and Fulv+ACSI) and overlapping for ACSS2, ER⍰, and H3K27ac (n=3 per group). **(E)** Changes in the magnitude of binding sites with each treatment (Veh, Fulv, ACSI, and Fulv+ACSI) for the top 10% binding sites of ACSS2 (left) and the top 1% binding sites of ER⍰ (right) (n=3 per group). **(F)** All binding sites for Fulv treatment of ER⍰, ACSS2, and H3K27 acetylation mapped on concatenated binding sites of Veh and Fulv treatments of ACSS2 pulldown to determine the changes with Fulv’s effect (left). The top 4 clusters with the overlapping binding for all three components were isolated, and the changes in the binding profile of these were compared across different treatments (Veh, Fulv, ACSI, and Fulv+ACSI) for ACSS2 (middle) and ER⍰ (right) pulldowns (n=3 per group).

| Treatment | Gene sets |
| --- | --- |
| Veh | • Estrogen Response (Early & Late)<br>• Glycolysis<br>• Myogenesis<br>• Mitosis<br>• Cell morphogenesis<br>• Nucleotide binding<br>• Transcription regulator activity<br>• Molecular function regulator activity |
| Fulv | • Early estrogen response<br>• mTORC1 signaling<br>• G2M checkpoint<br>• Mitosis<br>• Fatty acid metabolism<br>• RNA polymerase II transcription<br>• ESR mediated signaling<br>• Nuclear receptor signaling<br>• Estrogen dependent gene expression<br>• Nucleotide binding<br>• Cell cycle process<br>• Intracellular signal transduction<br>• Cellular response to stress<br>• Growth |
| ACSI | • Cell morphogenesis<br>• Cell differentiation regulation<br>• Intracellular transport<br>• Cellular response to stress<br>• Hormone response<br>• Transcription factor binding |
| Comb | • Transport regulation<br>• Phospholipid binding<br>• Phosphatidylinositol binding |

Motif analysis revealed that Fulv treatment enriched for transcription factors involved in DNA replication (E2F4 for ACSS2), estrogen response elements (ERE for ERα), and oncogenic transcription factors (Elk1 for H3K27ac) (**Suppl. Fig. 3A-C**). Notably, ERE enrichment persisted with combination treatment for ERα binding (**Suppl. Fig. 3B**), while other treatment groups showed distinct motif profiles with most motifs being treatment-specific (**Suppl. Fig. 3D**). Motifs common to all three factors in Veh and Fulv groups were associated with cell proliferation and oncogene expression, but these enrichments were lost with combination treatment (**Suppl. Fig. 3E**).

Genomic distribution analysis showed that ERα-enriched regions were predominantly located in promoters, with levels decreasing under combination treatment. H3K27ac was similarly concentrated in promoter regions with relatively stable distribution across treatments. In contrast, ACSS2 showed dramatically altered binding patterns: while Veh and Fulv treatments showed predominant promoter localization followed by distal intergenic regions, combination treatment shifted binding primarily to distal intergenic regions followed by introns (**Suppl. Fig. 4**). This substantial redistribution likely reflects the overall loss of ACSS2 binding sites with combination treatment.

Given that ACSS2 enrichment at chromatin increases with Fulv treatment, to investigate these genes and their roles in cancer progression, we determined the co-occupied binding sites of ACSS2 and ERα, along with H3K27 acetylation (**Fig. 5F-left**). The first four clusters in the heatmap represent genes with binding sites for both ACSS2 and ERα, as well as H3K27 acetylation. We then focused on these four clusters and examined how different treatments affected gene binding within these clusters. We observed that Fulv treatment substantially increased binding, whereas the ACSS2 inhibitor combination resulted in an almost complete loss of binding at these sites for ACSS2 (**Fig. 5F-middle**). For ERα binding (**Fig. 5F-right**) and H3K27 acetylation (**Suppl. Fig. 5A-left**), we also observed an increase with Fulv and a decrease with combination treatments. GSEA analysis of the genes in these clusters revealed they were almost exclusively involved in cell proliferation (**Suppl. Fig. 5B**). We also demonstrated that among all ACSS2-ERα-H3K27ac enriched regions, the majority were located in promoter regions, and the motifs in these regions were enriched for zinc finger proteins (**Suppl. Fig. 6**). Investigation of cluster 5 in H3K27ac pulldown (**Fig. 5C-right**) showed increased histone acetylation with Fulv despite unchanged chromatin availability. Although few genes in this cluster bind to ACSS2 or ERα, those that were present showed increased binding levels of both ACSS2 and ERα with Fulv and decreased levels with combination treatment (**Suppl. Fig. 7**).

Examining changes in binding levels of two histone acetylase genes (KAT5 and KAT6B) showed increased chromatin enrichment for all three factors (ACSS2 and ER⍰ bindings and H3K27 acetylation) with Fulv and decreased levels with combination treatment. We observed the same pattern for a transcription factor (SP1), a cell cycle regulation gene (CCND1), and genes crucial in cell metabolism (IGF1R and NAMPT), indicating that Fulv significantly alters cell metabolism, potentially contributing to therapy resistance (**Suppl. Fig. 8**).

These findings demonstrate that ACSS2 associates with chromatin, forming complexes that correlate with ERα binding and H3K27 acetylation. The co-localization of these factors at promoter regions following Fulv treatment, and their disruption by ACSS2 inhibition, suggests a coordinated metabolic-epigenetic regulatory mechanism underlying endocrine resistance.

### ACSS2 Inhibition Reverses Fulv-Induced Transcriptional Reprogramming and Reduces Metastatic Burden

To validate the therapeutic potential of ACSS2 inhibition, we tested the combination of ACSI and Fulv in an in vivo metastasis model using tail vain delivery of MCF7-ESR1^Y537S^ cells, which exclusively results in liver metastatic tumors^14,15,31^. At treatment initiation, most mice had already developed tumors of significant size. By the end of treatment, all groups showed continued tumor growth at varying levels (**Fig. 6A**). However, mice receiving combination Fulv + ACSI treatment showed significantly reduced increases in metastatic burden compared to the start of the treatment (**Fig. 6B and 6C**). Additionally, combination-treated mice maintained weight (**Fig. 6D**) and exhibited increased food consumption (**Suppl. Fig. 9**), indicating improved overall health and appetite.

**Figure 6.**
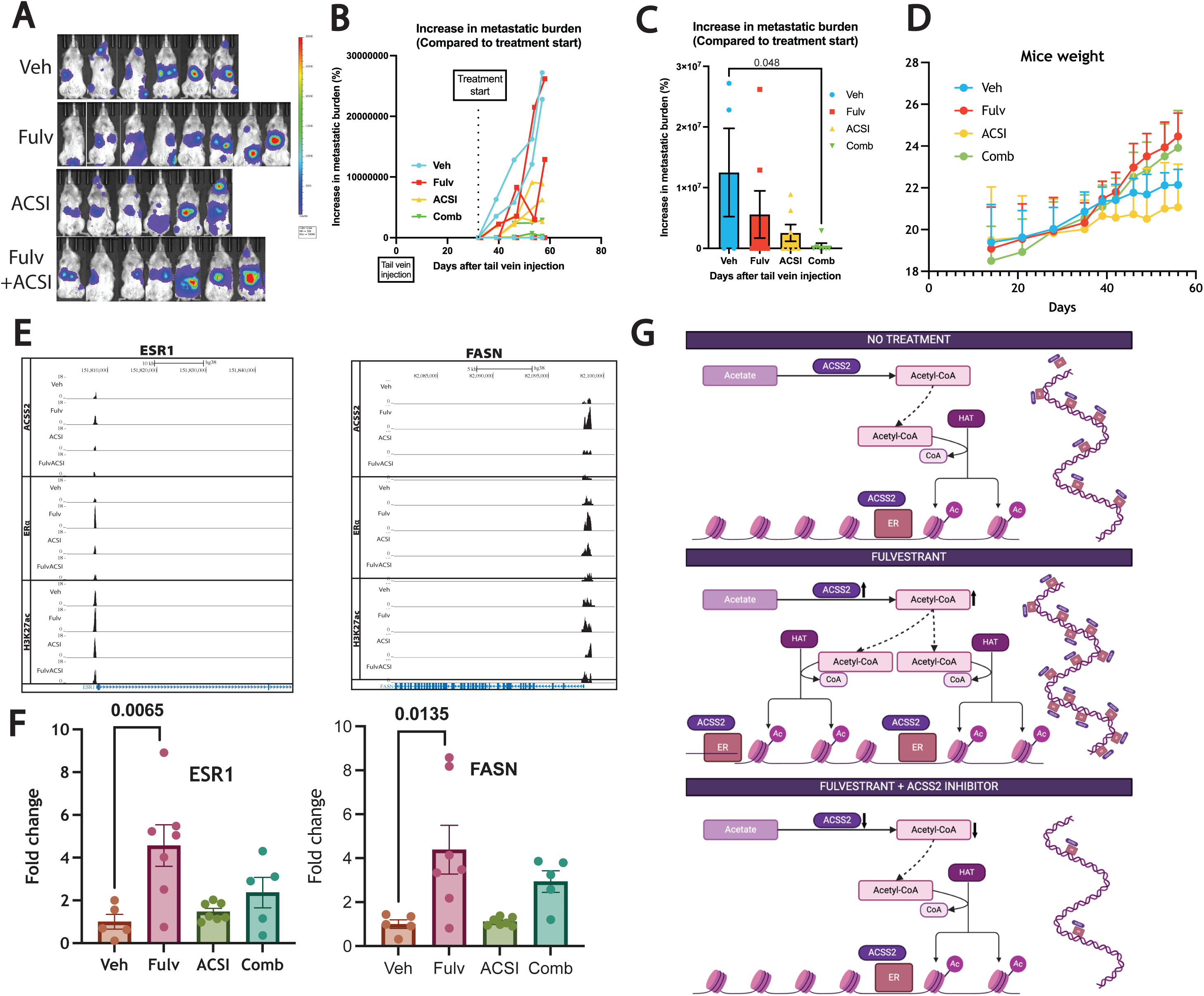
ACSS2 Inhibition Reverses Fulv-Induced Transcriptional Reprogramming and Reduces Metastatic Burden In Vivo. **(A)** IVIS images of mice treated with Veh (N=6), Fulv (N=8), ACSI (N=6), or Fulv+ACSI Comb (N=7) at the end of the treatments (Day 56). Line graph depicting the timeline change of the metastatic burden starting with the day of Fulv treatment start **(B)**, bar graph showing the increase in metastatic burden compared to treatment start (Day 32) at the end of the treatment (the day before sacrifice) (Day 56) **(C)** and the changes in the mice weight between different groups **(D)**. **(E)** Changes in the binding intensities of ESR1 and FASN gene with different treatments (Veh, Fulv, ACSI, Comb) for ACSS2 and ER⍰ pulldowns, H3K27 acetylation (n=3 per group). **(F)** Bar graph showing the fold changes between Veh (N=5), Fulv (N=7), ACSI (N=7), and Comb (N=5) treatments for the gene expression of ESR1 and FASN genes obtained from qPCR. **(G)** Proposed model for the role of ACSS2 in epigenetic regulation associated with Fulv resistance.

To confirm that the in vivo therapeutic effects correlate with our mechanistic findings, we examined the expression of genes identified as ACSS2-ERα-H3K27ac targets in xenograft tumor samples from treated mice. We examined expression of ESR1 and FASN, which has ACSS2-ERα-H3K27ac binding, which was reversed by combination therapy in MCF7-ESR1^Y537S^ cells (**Fig. 6E**). Quantitative PCR analysis of tumors from the animal study confirmed that the gene expression profiles (**Fig. 6F**) aligned with our CUT&RUN findings (**Fig. 6E**), demonstrating that the chromatin-level mechanisms observed in vitro translate to the in vivo setting and contribute to the therapeutic efficacy of the combination approach.

## 5. Discussion

Our study reveals a previously unrecognized nuclear role for ACSS2 in coordinating endocrine resistance through metabolic and epigenetic mechanisms. Our clinical analysis of 192 breast cancer patients revealed that liver metastasis represents a particularly aggressive form of metastatic disease, with significantly reduced survival rates and poor response to Fulv therapy. These findings align with our previous observations ^21^ and emerging clinical evidence that liver metastases create unique therapeutic challenges, potentially due to the metabolically active hepatic microenvironment that may facilitate adaptive resistance mechanisms. The disparity in Fulv efficacy between liver and non-liver metastatic sites underscores the urgent need for novel therapeutic strategies specifically targeting the metabolic vulnerabilities that drive resistance in this challenging patient population.

While ACSS2 can be located in both the cytosol and the nucleus ^15^, and its cytoplasmic functions in fatty acid synthesis are well-established, we demonstrated that Fulv treatment promotes ACSS2 nuclear translocation in therapy-resistant ESR1-mutant breast cancer cells. This nuclear ACSS2 forms functional chromatin complexes with ERα and drives H3K27 acetylation at promoters of proliferation and metabolic genes. Importantly, this represents a fundamental shift from viewing ACSS2 primarily as a metabolic enzyme to recognizing its direct role in transcriptional regulation by ERα and endocrine resistance.

Our findings are consistent with previous studies, which reported a key role for ACSS2 in tumor progression. It is shown that under lipid-depleted, low oxygen conditions, ACSS2 expression is significantly upregulated, and when silenced, it hinders tumor growth in breast cancer tumors ^32^. Similar tumor growth suppression was observed in triple-negative breast tumors with decreased ACSS2 expression ^22^, as well as myeloma ^33^ and liver cancer ^17^, where ACSS2 overexpression facilitated disease progression while inhibition decreased cell growth. However, our work extends these findings by demonstrating that ACSS2 functions beyond cellular metabolism to directly regulate chromatin states and gene expression. The discovery that ACSS2 inhibition disrupts these chromatin complexes and reverses Fulv-induced gene expression changes provides mechanistic insight into how metabolic enzymes can directly influence epigenetic landscapes. This metabolic-epigenetic axis represents a novel therapeutic vulnerability, as targeting ACSS2 simultaneously disrupts both altered cellular metabolism and the transcriptional programs that maintain the resistant phenotype.

Our multi-modal experimental approach provides convergent evidence for the ACSS2-mediated resistance mechanism. PDX models demonstrated that Fulv-resistant liver metastatic tumors has increased fatty acid metabolism upon Fulv treatment. Isotope tracing experiments confirmed that Fulv treatment redirects acetate flux toward lipid synthesis, consistent with previous reports showing that under stressful conditions, cancer cells predominantly utilize acetate as a source of acetyl-CoA for fatty acid synthesis ^34,35^. Critically, our CUT&RUN analysis revealed the chromatin-level consequences of this metabolic reprogramming. Previous work has shown that ACSS2 works with transcription factors to provide acetyl groups for histone acetylation, leading to gene activation ^16,36^. Building on these findings, we demonstrate that the ACSS2-ERα complex promotes cell proliferation by activating specific genes through H3K27 acetylation. This is particularly significant given that histone H3 acetylation at lysine 27 (H3K27ac) serves as a key marker distinguishing active enhancers from inactive ones, and ESR1 gene transcription is specifically regulated by H3K27ac signaling ^37^. Additionally, H3K27ac levels increase at ER-controlled sites upon estradiol (E2) treatment ^38^.

Our RNA sequencing demonstrated that ACSS2 inhibition reverses Fulv-induced transcriptional changes, while in vivo studies validated the therapeutic potential of combination therapy. The convergence of these findings supports a model where Fulv treatment inadvertently promotes a metabolic-epigenetic adaptation: increased nuclear ACSS2 provides local acetyl-CoA for histone acetylation, driving expression of genes that promote cell survival and proliferation (**Fig. 6G**). This mechanism may be particularly pronounced in liver metastases due to the abundant acetate availability in the hepatic environment.

The identification of ACSS2 as a driver of Fulv resistance provides several therapeutic opportunities. First, ACSS2 inhibitors could be developed as combination agents with existing endocrine therapies, potentially improving outcomes for patients with liver metastases who currently have limited treatment options. Our previous research showed that elevated NAMPT expression contributes to therapy resistance, and its inhibition significantly reduces tumor burden and cell viability ^15^. The current findings suggest that ACSS2-mediated epigenetic regulation of NAMPT expression might represent an important link between these mechanism facilitating therapy resistance. Second, ACSS2 expression or activity could serve as a biomarker to identify patients most likely to benefit from combination approaches. Our findings also suggest that targeting metabolic-epigenetic crosstalk represents a promising therapeutic strategy beyond breast cancer. The concept that metabolic enzymes can directly regulate chromatin states opens new avenues for drug development, particularly in cancers characterized by metabolic reprogramming and epigenetic dysregulation.

There are several limitations in our study that warrant consideration. Our in vivo studies used relatively low doses and short treatment durations due to drug solubility constraints, which may have limited therapeutic efficacy. In these studies, we opted for the tumors to fully establish, which led to a late start to treatment. Additionally, the limitation of the ACSI we used was low solubility, which did not allow us to use the drugs at the concentrations we initially planned (150 mg/kg/day). Despite the low dose and short treatment period, we observed that the combination treatment inhibited increase in metastatic burden, and increased dosage and longer treatment periods could provide more effective decrease in metastatic burden and improved symptoms. Future studies should optimize dosing regimens and explore alternative ACSS2 inhibitors with improved pharmacological properties.

## 6. Conclusion

Our work establishes a new paradigm for understanding endocrine resistance that integrates metabolic reprogramming with epigenetic regulation. The dependency on histone acetylation, particularly H3K27ac, represents a critical mechanism in cancer cell survival and proliferation and may provide a therapeutic vulnerability that could be exploited in cancers heavily reliant on these pathways. Future research should explore whether similar metabolic-epigenetic mechanisms operate in other forms of therapy resistance and whether targeting these pathways could improve outcomes across multiple cancer types. Clinical translation will require the development of optimized ACSS2 inhibitors, the identification of biomarkers to select appropriate patients, and the careful design of combination therapy regimens. Given the vulnerability of liver metastatic disease, future clinical trials should prioritize this patient population for early-phase studies.

**Supplementary Figure 1. (A)** Prevalence of metabolic diseases in patients with liver metastases (left) and non-liver metastases (right), before and after metastasis diagnosis. Abbreviations: HT, hypertension; DM, diabetes mellitus; CKD, chronic kidney disease; IHD, ischemic heart disease. **(B)** The changes in tumor volume with and without Fulv treatment for HCI013 (N=5 per group) PDX tumors.

**Supplementary Figure 2.** Heatmaps showing ATAC-Sequencing results for **(A)** all ATAC-seq data, **(B)** overlapping sites (for ER⍰, ACSS2, and H3K27ac), **(C)** ACSS2, **(D)** ER⍰, and **(E)** H3K27ac (n=3 per group).

**Supplementary Figure 3.** Motif analysis for **(A)** ACSS2, **(B)** ER⍰, **(C)** H3K27ac pulldowns under different treatments (Veh, Fulv, ACSI, and Fulv+ACSI). **(D)** Venn diagrams showing the number of motifs exclusive to one treatment or overlapping with different treatments (Veh, Fulv, ACSI, and Fulv+ACSI) for ACSS2, ER⍰, and H3K27ac (n=3 per group)). **(E)** Venn diagrams showing the number of motifs overlapping for ACSS2, ER⍰ bindings and H3K27ac. The analysis of some treatments (ER⍰ pulldown of ACSI and H3K27 acetylation of ACSI) did not yield any motifs, and those portions of the figure are left empty.

**Supplementary Figure 4.** Percentage of binding site distribution with different treatments (Veh, Fulv, ACSI, and Fulv+ACSI) for ACSS2, ER⍰, and H3K27ac (n=3 per group) pulldowns.

**Supplementary Figure 5. (A)** The top 4 clusters with the overlapping binding for ER⍰ and ACSS2 pulldowns and H3K27 acetylation from Figure 5F-left are isolated, and the changes in the binding profile of these with different treatments (Veh, Fulv, ACSI, and Fulv+ACSI) for H3K27 acetylation **(left)**, and ATAC-seq **(right)** results were shown in heatmaps. **(B)** GSEA results of the gene sets found in the top 4 clusters from Figure 5F-left.

**Supplementary Figure 6.** Motif analysis and binding site distribution of overlapping binding sites (n=3).

**Supplementary Figure 7. (A)** Heatmap for H3K27ac pulldown with different treatments (Veh, Fulv, ACSI, and Comb). Heatmaps for the changes in Cluster 5 of H3K27ac pulldown for **(B)** ATAC-seq**, (C)** ACSS2, **(D)** ER⍰, and **(E)** H3K27ac pulldowns (n=3 per group).

**Supplementary Figure 8.** Changes in the binding intensities of histone acetylases KAT5 and KAT6B; CCND1, SP1, and metabolic proteins IGF1R and NAMPT with different treatments (Veh, Fulv, ACSI, and Fulv+ACSI) (n=3 per group).

**Supplementary Figure 9.** The weight **(top)** and food consumption **(bottom)** levels of different treatments (Veh (N=7), Fulv (N=8), ACSI (N=7), or Fulv+ACSI Comb (N=8)) for the in vivo study.

## Notes

**Funding:** The research reported in this publication was supported by the Cancer Scholars for Translational and Applied Research (CSTAR) Program sponsored by the Cancer Center at Illinois and the Carle Cancer Center under Award Number CST EP082021 (to ANM) and by the National Institute of Biomedical Imaging and Bioengineering of the National Institutes of Health under Award Number T32EB019944 (to JYY and CPS). Further grant support is provided by University of Illinois, Office of the Vice Chancellor for Research, Campus Research Board grant RB23080, National Institute of Food and Agriculture, U.S. Department of Agriculture, award ILLU-698-924 and ILLU-698-331, Prairie Dragon Paddlers Award and Sylvia D. Stroup Scholar Award (to ZME) and by Department of Defense Era of Hope Scholar Award BC200206/W81XWH-20-BCRP-EOHS (to ERN).

### Competing Interest Statement

The authors have declared no competing interest.

