## Supplementary figures and images for "ACSS2-Mediated Metabolic-Epigenetic Crosstalk Drives Fulvestrant Resistance and Represents a Novel Therapeutic Target"

### Supplemental Figure 1

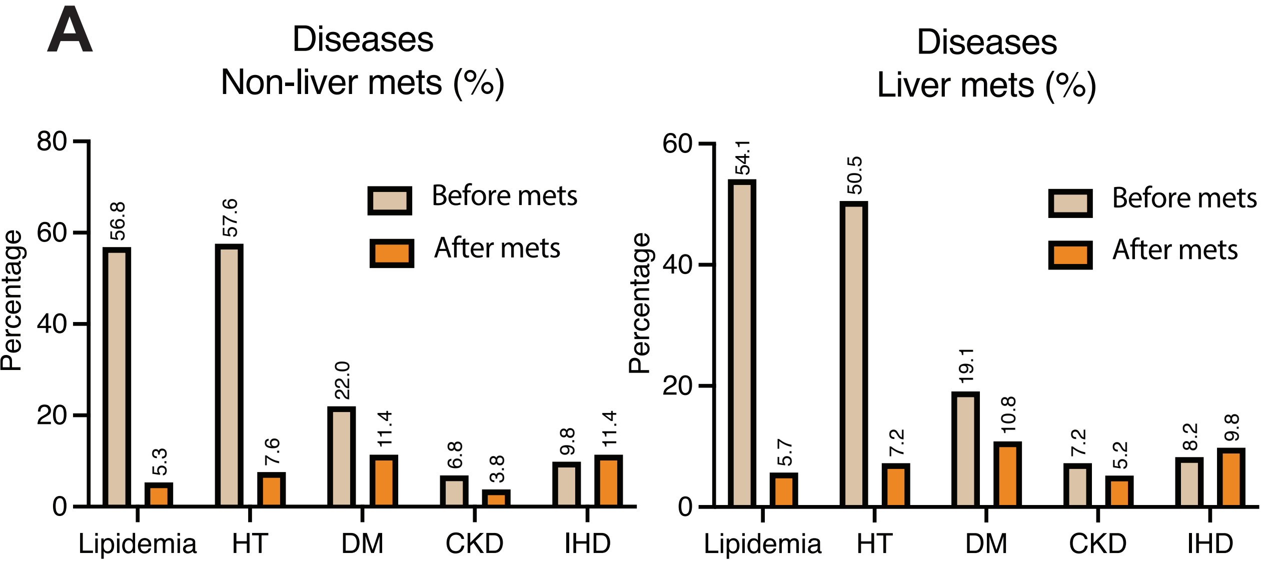


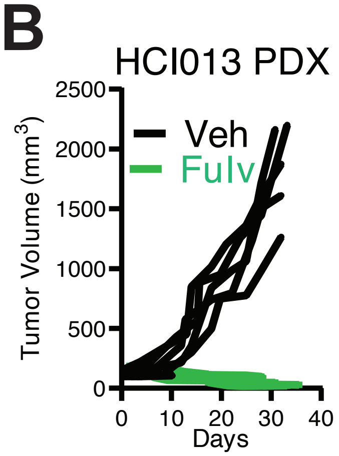


**Supplementary Figure 1**

### Supplemental Figure 2

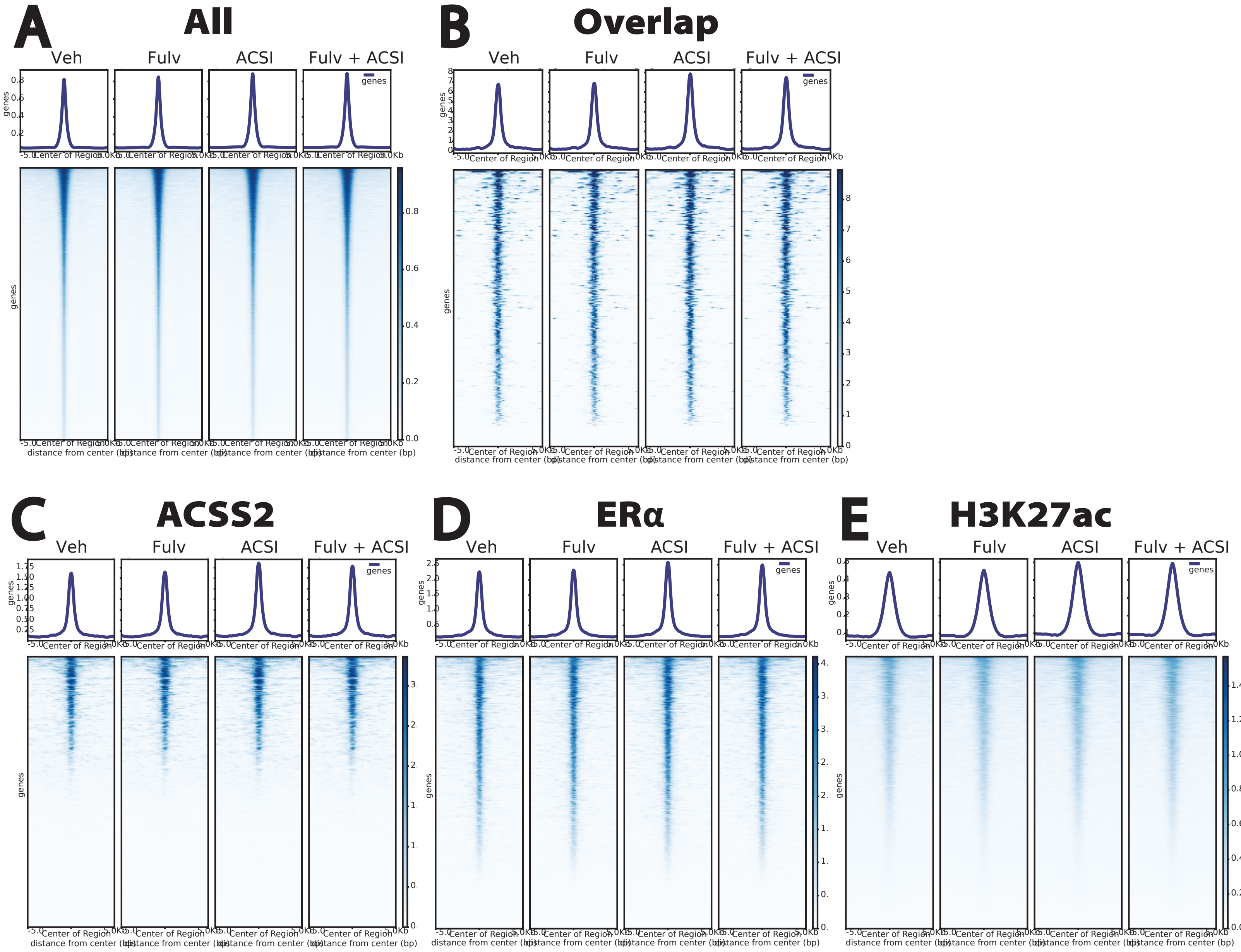


#### Supplementary Figure 2

### Supplemental Figure 4

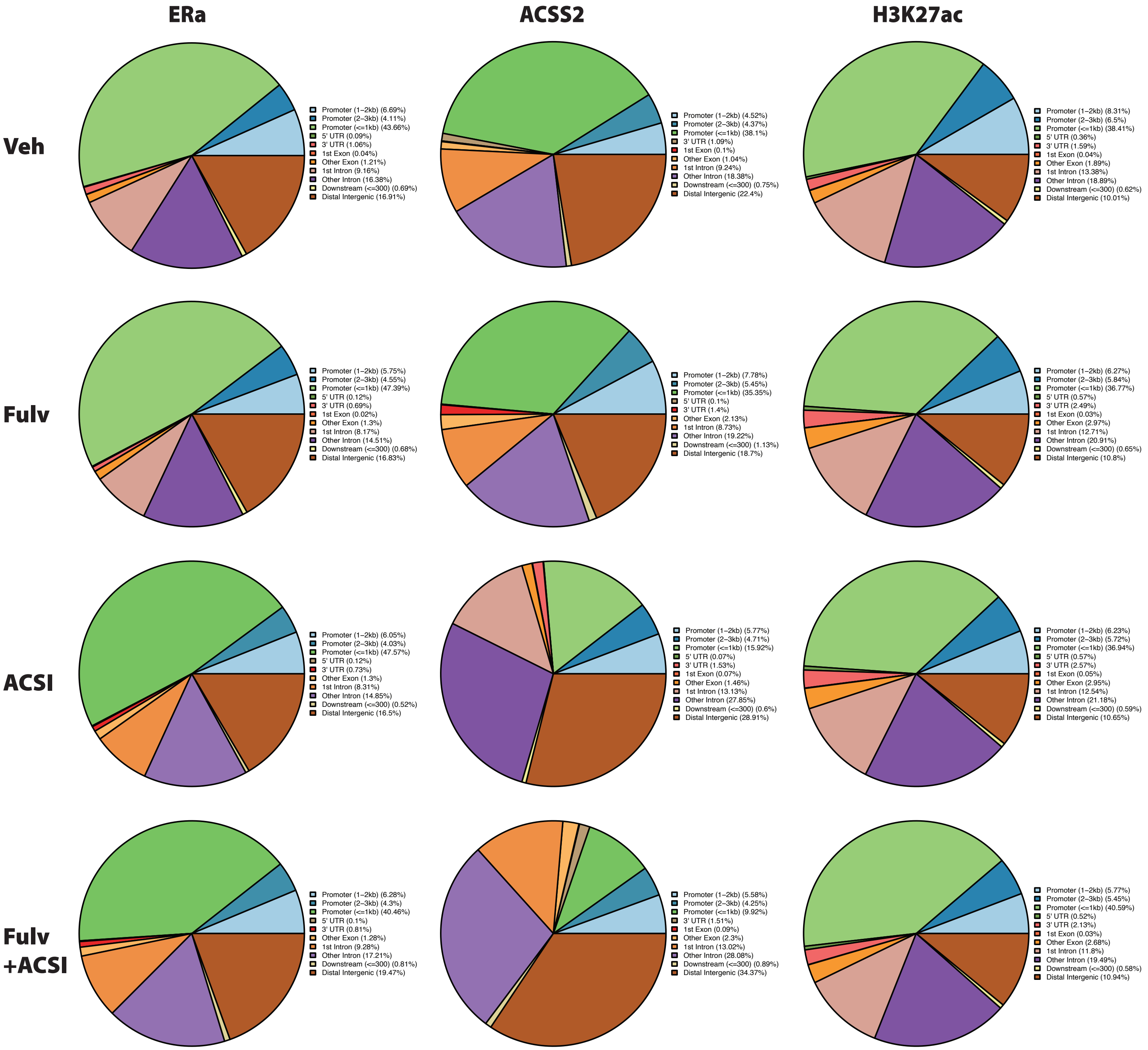


#### Supplementary Figure 4

### Supplemental Figure 5

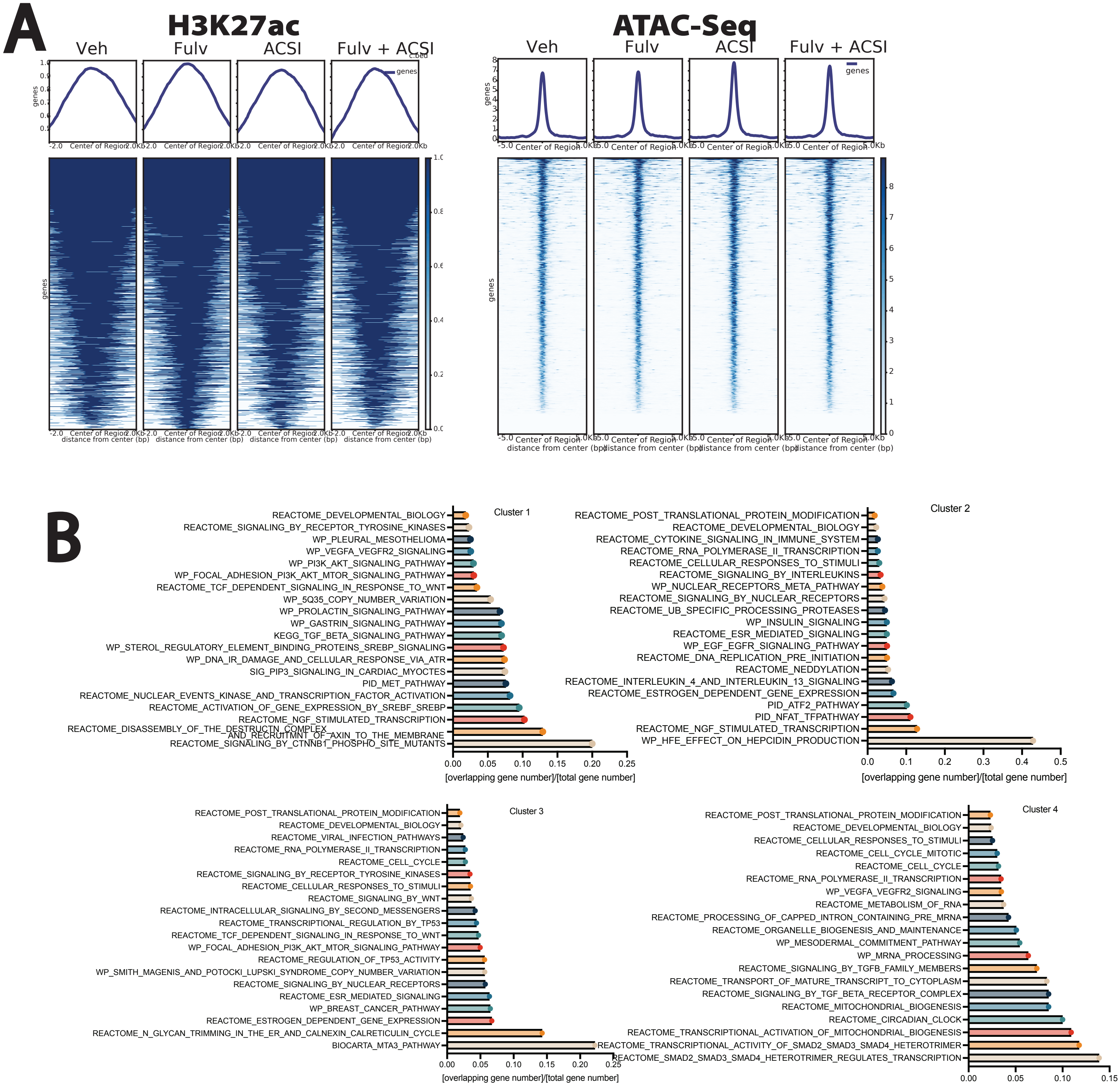


#### Supplementary Figure 5

### Supplemental Figure 7

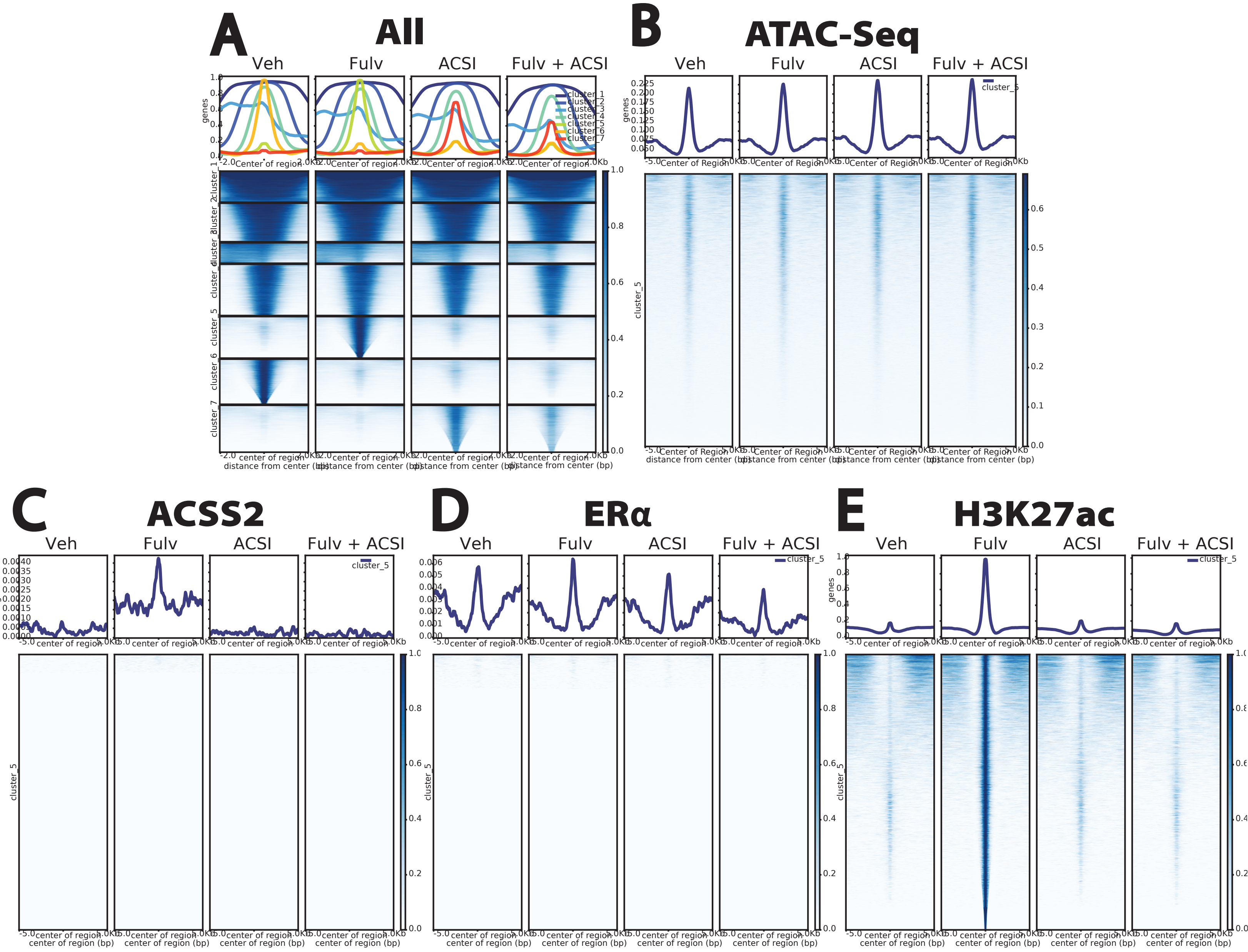


#### Supplementary Figure 7

### Supplemental Figure 8

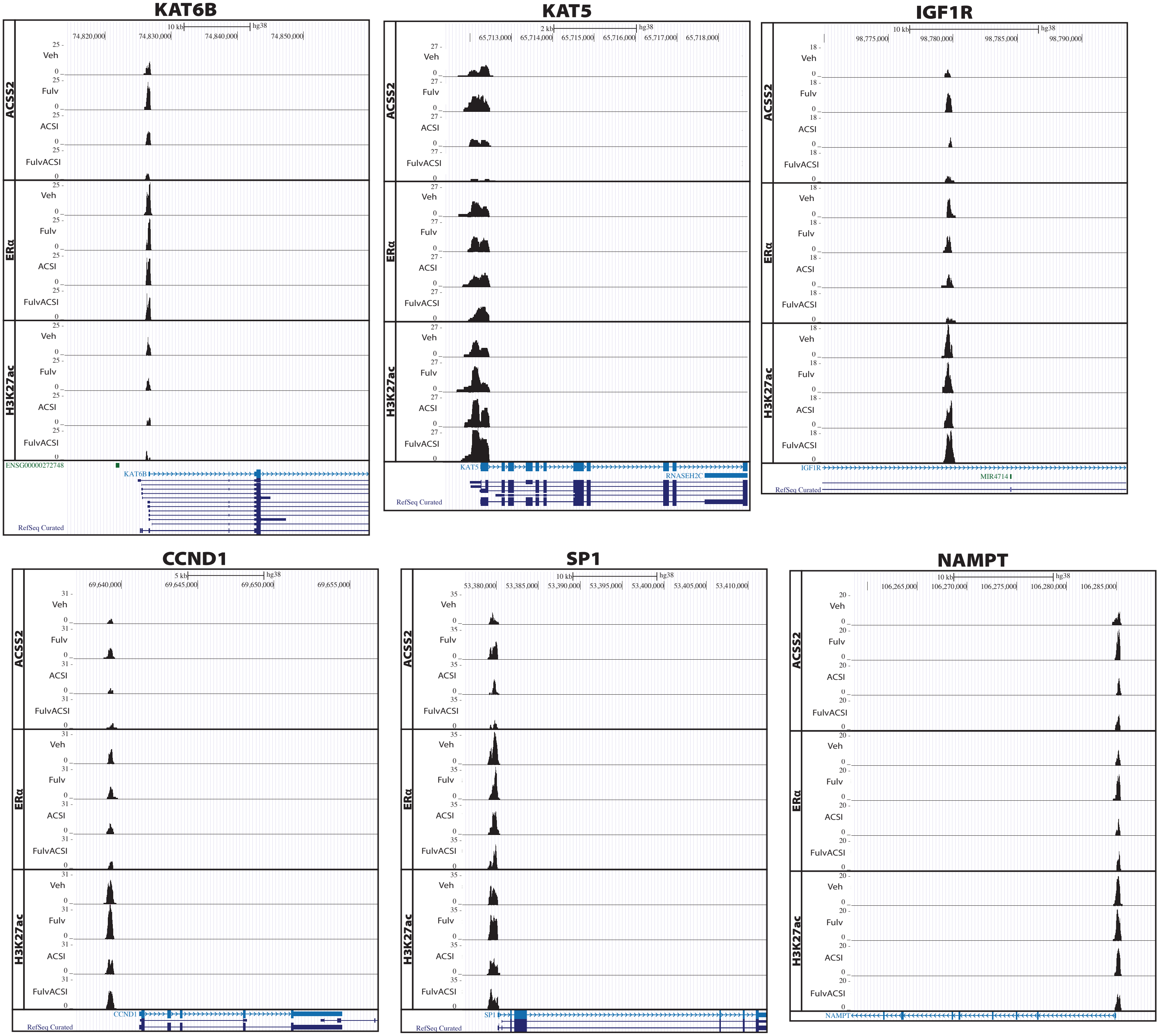


#### Supplementary Figure 8

### Supplemental Figure 9

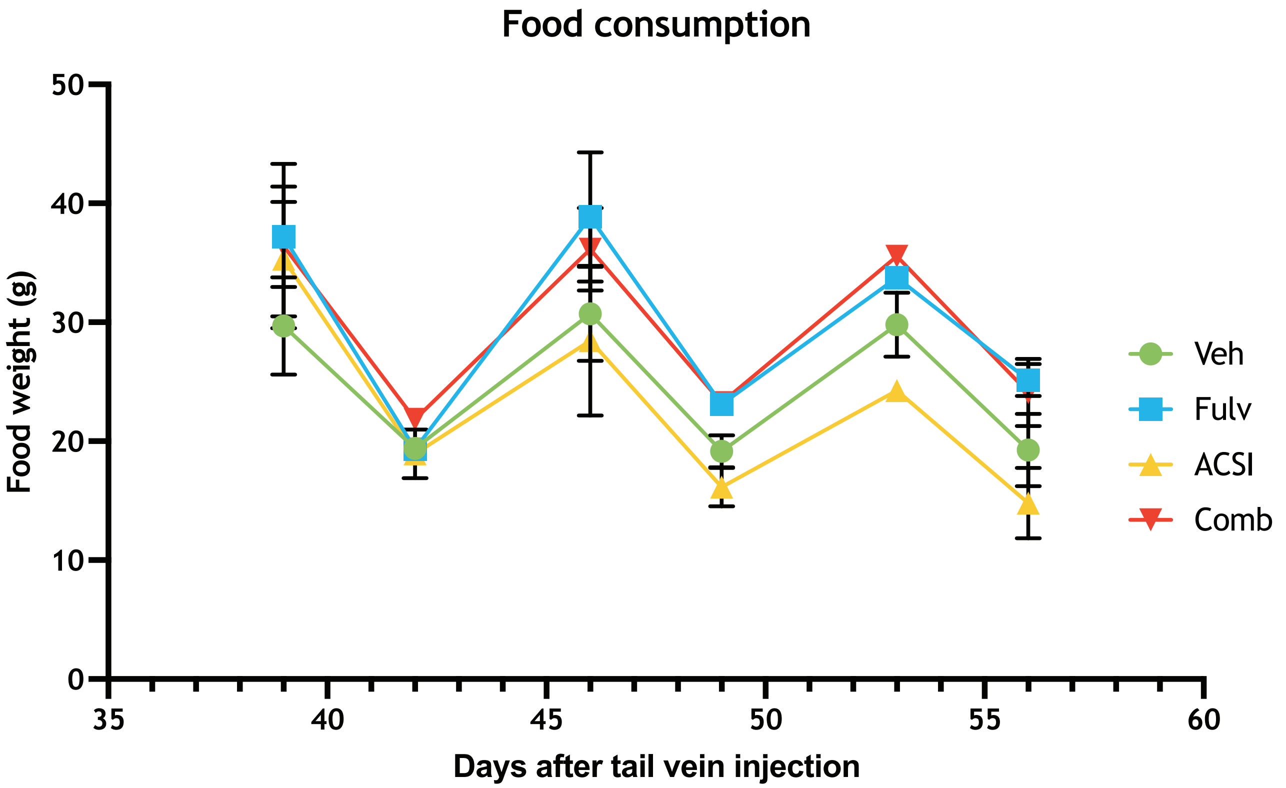


#### Supplementary Figure 9
