## Supplemental Figure 3 for "ACSS2-Mediated Metabolic-Epigenetic Crosstalk Drives Fulvestrant Resistance and Represents a Novel Therapeutic Target"

# A

#### ACSS2

##### Veh

| Rank | Motif | Name | P-value | % of Targets Sequences with Motif | % of Background Sequences with Motif |
|---|---|---|---|---|---|
| 1 | | KLF1(Zf)/HUDEP2-KLF1-CutnRun(GSE136251)/Homer | 1e-7 | 5.91% | 3.03% |
| 2 | | Sp2(Zf)/HEK293-Sp2.eGFP-ChIP-Seq(Encode)/Homer | 1e-6 | 8.87% | 5.37% |
| 3 | | NFY(CCAAT)/Promoter/Homer | 1e-6 | 3.18% | 1.31% |
| 4 | | Sp1(Zf)/Promoter/Homer | 1e-5 | 2.88% | 1.19% |
| 5 | | Ronin(THAP)/ES-Thap11-ChIP-Seq(GSE51522)/Homer | 1e-5 | 0.38% | 0.02% |
| 6 | | GFY(?)/Promoter/Homer | 1e-5 | 0.61% | 0.07% |
| 7 | | GFY-Staf(?,Zf)/Promoter/Homer | 1e-5 | 0.45% | 0.03% |
| 8 | | KLF14(Zf)/HEK293-KLF14.GFP-ChIP-Seq(GSE58341)/Homer | 1e-4 | 9.17% | 6.26% |
| 9 | | KLF5(Zf)/LoVo-KLF5-ChIP-Seq(GSE49402)/Homer | 1e-4 | 6.90% | 4.39% |
| 10 | | KLF6(Zf)/PDAC-KLF6-ChIP-Seq(GSE64557)/Homer | 1e-4 | 5.69% | 3.49% |

##### Fulv

| Rank | Motif | Name | P-value | % of Targets Sequences with Motif | % of Background Sequences with Motif |
|---|---|---|---|---|---|
| 1 | | Sp5(Zf)/mES-Sp5.Flag-ChIP- | 1e-16 | 6.78% | 3.80% |
| 2 | | Sp1(Zf)/Promoter/Homer | 1e-15 | 3.05% | 1.29% |
| 3 | | KLF1(Zf)/HUDEP2-KLF1-CutnRun(GSE136251)/Homer | 1e-13 | 5.96% | 3.43% |
| 4 | | NFY(CCAAT)/Promoter/Homer | 1e-12 | 2.91% | 1.30% |
| 5 | | Sp2(Zf)/HEK293-Sp2.eGFP-ChIP-Seq(Encode)/Homer | 1e-11 | 9.20% | 6.22% |
| 6 | | KLF3(Zf)/MEF-Klf3-ChIP-Seq(GSE44748)/Homer | 1e-10 | 3.08% | 1.54% |
| 7 | | KLF14(Zf)/HEK293-KLF14.GFP-ChIP-Seq(GSE58341)/Homer | 1e-9 | 9.83% | 7.00% |
| 8 | | KLF5(Zf)/LoVo-KLF5-ChIP-Seq(GSE49402)/Homer | 1e-9 | 7.52% | 5.09% |
| 9 | | E2F4(E2F)/K562-E2F4-ChIP-Seq(GSE31477)/Homer | 1e-9 | 3.05% | 1.60% |

##### ACSI

| Rank | Motif | Name | P-value | % of Targets Sequences with Motif | % of Background Sequences with Motif |
|---|---|---|---|---|---|
| 1 | | WRKY47(WRKY)/colamp-WRKY47-DAP-Seq(GSE60143)/Homer | 1e-7 | 1.02% | 0.13% |
| 2 | | GATA4(C2C2gata)/col-GATA4-DAP-Seq(GSE60143)/Homer | 1e-6 | 1.92% | 0.45% |
| 3 | | Hoxd12(Homeobox)/ChickenMSG-Hoxd12.Flag-ChIP-Seq(GSE86088)/Homer | 1e-6 | 7.24% | 3.66% |
| 4 | | Unknown4/Drosophila-Promoters/Homer | 1e-6 | 0.90% | 0.09% |
| 5 | | HOXA9(Homeobox)/HSC-Hoxa9-ChIP-Seq(GSE33509)/Homer | 1e-5 | 2.26% | 0.70% |
| 6 | | TATA-box/SacCer-Promoters/Homer | 1e-5 | 1.92% | 0.50% |
| 7 | | AT3G42860(zfGRF)/col-AT3G42860- | 1e-5 | 0.79% | 0.13% |
| 8 | | Jun-AP1(bZIP)/K562-cJun-ChIP-Seq(GSE31477)/Homer | 1e-4 | 1.02% | 0.19% |
| 9 | | Unknown6/Drosophila-Promoters/Homer | 1e-4 | 5.43% | 2.86% |
| 10 | | Sox6(HMG)/Myotubes-Sox6-ChIP-Seq(GSE32627)/Homer | 1e-4 | 5.43% | 2.94% |

##### Fulv ACSI

| Rank | Motif | Name | P-value | % of Targets Sequences with Motif | % of Background Sequences with Motif |
|---|---|---|---|---|---|
| 1 | | GTL1(Trihelix)/colamp-GTL1-DAP-Seq(GSE60143)/Homer | 1e-2 | 6.77% | 4.11% |
| 2 | | AT1G24250(Orphan)/col-AT1G24250-DAP-Seq(GSE60143)/Homer | 1e-2 | 1.65% | 0.56% |
| 3 | | MYB65(MYB)/colamp-MYB65-DAP-Seq(GSE60143)/Homer | 1e-2 | 5.12% | 3.05% |
| 4 | | SND2(NAC)/colamp-SND2-DAP-Seq(GSE60143)/Homer | 1e-2 | 2.15% | 0.92% |
| 5 | | Zfp57(Zf)/H1-ZFP57.HA-ChIP-Seq(GSE115387)/Homer | 1e-2 | 0.99% | 0.26% |
| 6 | | AT5G47660(Trihelix)/colamp-AT5G47660-DAP-Seq(GSE60143)/Homer | 1e-2 | 8.09% | 5.60% |
| 7 | | STOP1(C2H2)/colamp-STOP1-DAP-Seq(GSE60143)/Homer | 1e-2 | 1.65% | 0.67% |
| 8 | | RORgt(NR)/EL4-RORgt.Flag-ChIP-Seq(GSE56019)/Homer | 1e-2 | 0.99% | 0.30% |
| 9 | | RORgt(NR)/EL4-RORgt.Flag-ChIP-Seq(GSE56019)/Homer | 1e-2 | 0.99% | 0.30% |

# B

#### ERα

##### Veh

| Rank | Motif | Name | P-value | % of Targets Sequences with Motif | % of Background Sequences with Motif |
|---|---|---|---|---|---|
| 1 | | NFY(CCAAT)/Promoter/Homer | 1e-36 | 3.69% | 1.35% |
| 2 | | Seq(Encode)/Homer | 35 | 8.87% | 4.92% |
| 3 | | Sp5(Zf)/mES-Sp5.Flag-ChIP-Seq(GSE72989)/Homer | 1e-34 | 6.24% | 3.05% |
| 4 | | Sp1(Zf)/Promoter/Homer | 1e-33 | 2.95% | 0.98% |
| 5 | | KLF1(Zf)/HUDEP2-KLF1-CutnRun(GSE136251)/Homer | 1e-31 | 5.49% | 2.66% |
| 6 | | KLF3(Zf)/MEF-Klf3-ChIP-Seq(GSE44748)/Homer | 1e-27 | 3.02% | 1.17% |
| 7 | | KLF5(Zf)/LoVo-KLF5-ChIP-Seq(GSE49402)/Homer | 1e-23 | 6.90% | 4.00% |
| 8 | | KLF6(Zf)/PDAC-KLF6-ChIP-Seq(GSE64557)/Homer | 1e-20 | 5.48% | 3.12% |
| 9 | | KLF14(Zf)/HEK293-KLF14.GFP-ChIP-Seq(GSE58341)/Homer | 1e-19 | 8.56% | 5.62% |
| 10 | | FOXA1(Forkhead)/MCF7-FOXA1-ChIP-Seq(GSE26831)/Homer | 1e-15 | 3.36% | 1.80% |

##### Fulv

| Rank | Motif | Name | P-value | % of Targets Sequences with Motif | % of Background Sequences with Motif |
|---|---|---|---|---|---|
| 1 | | KLF1(Zf)/HUDEP2-KLF1-CutnRun(GSE136251)/Homer | 1e-29 | 6.36% | 3.01% |
| 2 | | Sp1(Zf)/Promoter/Homer | 1e-25 | 3.28% | 1.18% |
| 3 | | KLF5(Zf)/LoVo-KLF5-ChIP-Seq(GSE49402)/Homer | 1e-24 | 8.02% | 4.47% |
| 4 | | Sp2(Zf)/HEK293-Sp2.eGFP-ChIP-Seq(Encode)/Homer | 1e-23 | 9.37% | 5.54% |
| 5 | | Sp5(Zf)/mES-Sp5.Flag-ChIP-Seq(GSE72989)/Homer | 1e-22 | 6.51% | 3.46% |
| 6 | | KLF3(Zf)/MEF-Klf3-ChIP-Seq(GSE44748)/Homer | 1e-19 | 3.14% | 1.30% |
| 7 | | KLF6(Zf)/PDAC-KLF6-ChIP-Seq(GSE64557)/Homer | 1e-18 | 6.45% | 3.66% |
| 8 | | NFY(CCAAT)/Promoter/Homer | 1e-18 | 2.90% | 1.20% |
| 9 | | ERE(NR),IR3/MCF7-ERa-ChIP-Seq(Unpublished)/Homer | 1e-15 | 1.30% | 0.36% |
| 10 | | KLF14(Zf)/HEK293-KLF14.GFP-ChIP-Seq(GSE58341)/Homer | 1e-15 | 9.61% | 6.45% |

##### ACSI

##### Fulv ACSI

| Rank | Motif | Name | P-value | % of Targets Sequences with Motif | % of Background Sequences with Motif |
|---|---|---|---|---|---|
| 1 | | NFY(CCAAT)/Promoter/Homer | 1e-22 | 3.88% | 1.28% |
| 2 | | Sp5(Zf)/mES-Sp5.Flag-ChIP-Seq(GSE72989)/Homer | 1e-22 | 6.27% | 2.74% |
| 3 | | Sp2(Zf)/HEK293-Sp2.eGFP-ChIP-Seq(Encode)/Homer | 1e-21 | 8.83% | 4.59% |
| 4 | | KLF1(Zf)/HUDEP2-KLF1-CutnRun(GSE136251)/Homer | 1e-17 | 5.38% | 2.45% |
| 5 | | KLF14(Zf)/HEK293-KLF14.GFP-ChIP-Seq(GSE58341)/Homer | 1e-14 | 8.76% | 5.17% |
| 6 | | Sp1(Zf)/Promoter/Homer | 1e-11 | 2.49% | 0.96% |
| 7 | | KLF5(Zf)/LoVo-KLF5-ChIP-Seq(GSE49402)/Homer | 1e-10 | 6.19% | 3.61% |
| 8 | | KLF3(Zf)/MEF-Klf3-ChIP-Seq(GSE44748)/Homer | 1e-10 | 2.60% | 1.09% |
| 9 | | ERE(NR),IR3/MCF7-ERa-ChIP-Seq(Unpublished)/Homer | 1e-7 | 1.10% | 0.33% |
| 10 | | Atf1(bZIP)/K562-ATF1-ChIP-Seq(GSE31477)/Homer | 1e-6 | 2.17% | 1.04% |

# C

## H3K27ac

**Veh**

| Rank | Motif | Name | P-value | % of Targets Sequences with Motif | % of Background Sequences with Motif |
|---|---|---|---|---|---|
| 1 | | Jun-AP1(bZIP)/K562-cJun-ChIP-Seq(GSE31477)/Homer | 1e-18 | 0.54% | 0.27% |
| 2 | | Fosl2(bZIP)/3T3L1-Fosl2-ChIP-Seq(GSE56872)/Homer | 1e-14 | 0.69% | 0.41% |
| 3 | | GRHL2(CP2)/HBE-GRHL2-ChIP-Seq(GSE46194)/Homer | 1e-14 | 0.79% | 0.49% |
| 4 | | E2FA(E2FDP)/colamp-E2FA-DAP-Seq(GSE60143)/Homer | 1e-13 | 0.72% | 0.44% |
| 5 | | FOXA1(Forkhead)/MCF7-FOXA1-ChIP-Seq(GSE26831)/Homer | 1e-12 | 2.99% | 2.39% |
| 6 | | KLF14(Zf)/HEK293-KLF14.GFP-ChIP-Seq(GSE58341)/Homer | 1e-12 | 4.92% | 4.15% |
| 7 | | KLF1(Zf)/HUDEP2-KLF1-CutnRun(GSE136251)/Homer | 1e-12 | 2.53% | 1.99% |
| 8 | | Fox:Ebox(Forkhead,bHLH)/Panc1-Foxa2-ChIP-Seq(GSE47459)/Homer | 1e-11 | 2.01% | 1.54% |
| 9 | | PHA-4(Forkhead)/cElegans-Embryos-PHA4-ChIP-Seq(modEncode)/Homer | 1e-11 | 9.28% | 8.28% |
| 10 | | Fra2(bZIP)/Striatum-Fra2-ChIP-Seq(GSE43429)/Homer | 1e-11 | 1.01% | 0.70% |

**Fulv**

| Rank | Motif | Name | P-value | % of Targets Sequences with Motif | % of Background Sequences with Motif |
|---|---|---|---|---|---|
| 1 | | GFY(?)/Promoter/Homer | 1e-18 | 0.19% | 0.05% |
| 2 | | Fli1(ETS)/CD8-FLI-ChIP-Seq(GSE20898)/Homer | 1e-14 | 2.81% | 2.16% |
| 3 | | CTCF(Zf)/CD4+-CTCF-ChIP-Seq(Barski_et_al.)/Homer | 1e-12 | 0.37% | 0.18% |
| 4 | | Jun-AP1(bZIP)/K562-cJun-ChIP-Seq(GSE31477)/Homer | 1e-11 | 0.52% | 0.29% |
| 5 | | Elk1(ETS)/Hela-Elk1-ChIP-Seq(GSE31477)/Homer | 1e-10 | 1.41% | 1.02% |
| 6 | | Sp2(Zf)/HEK293-Sp2.eGFP-ChIP-Seq(Encode)/Homer | 1e-10 | 4.49% | 3.78% |
| 7 | | Sp5(Zf)/mES-Sp5.Flag-ChIP-Seq(GSE72989)/Homer | 1e-10 | 2.86% | 2.30% |
| 8 | | JunB(bZIP)/DendriticCells-Junb-ChIP-Seq(GSE36099)/Homer | 1e-10 | 1.22% | 0.87% |
| 9 | | Sp1(Zf)/Promoter/Homer | 1e-10 | 0.99% | 0.68% |
| 10 | | Etv2(ETS)/ES-ER71-ChIP-Seq(GSE59402)/Homer | 1e-9 | 2.08% | 1.63% |

**ACSI**

**Fulv ACSI**

| Rank | Motif | Name | P-value | % of Targets Sequences with Motif | % of Background Sequences with Motif |
|---|---|---|---|---|---|
| 1 | | FOXA1(Forkhead)/MCF7-FOXA1-ChIP-Seq(GSE26831)/Homer | 1e-19 | 2.96% | 2.12% |
| 2 | | FOXA1(Forkhead)/LNCAP-FOXA1-ChIP-Seq(GSE27824)/Homer | 1e-19 | 3.57% | 2.64% |
| 3 | | FOXM1(Forkhead)/MCF7-FOXM1-ChIP-Seq(GSE72977)/Homer | 1e-17 | 2.95% | 2.15% |
| 4 | | ETS(ETS)/Promoter/Homer | 1e-15 | 0.89% | 0.51% |
| 5 | | LBD18(LOBAS2)/colamp-LBD18-DAP-Seq(GSE60143)/Homer | 1e-15 | 6.13% | 5.02% |
| 6 | | GABPA(ETS)/Jurkat-GABPa-ChIP-Seq(GSE17954)/Homer | 1e-14 | 2.37% | 1.71% |
| 7 | | Foxa2(Forkhead)/Liver-Foxa2-ChIP-Seq(GSE25694)/Homer | 1e-14 | 2.15% | 1.53% |
| 8 | | ETV4(ETS)/HepG2-ETV4-ChIP-Seq(ENCODE)/Homer | 1e-14 | 3.09% | 2.35% |
| 9 | | Foxo3(Forkhead)/U2OS-Foxo3-ChIP-Seq(E-MTAB-2701)/Homer | 1e-12 | 2.17% | 1.58% |
| 10 | | Foxa3(Forkhead)/Liver-Foxa3-ChIP-Seq(GSE77670)/Homer | 1e-11 | 0.85% | 0.52% |

# D

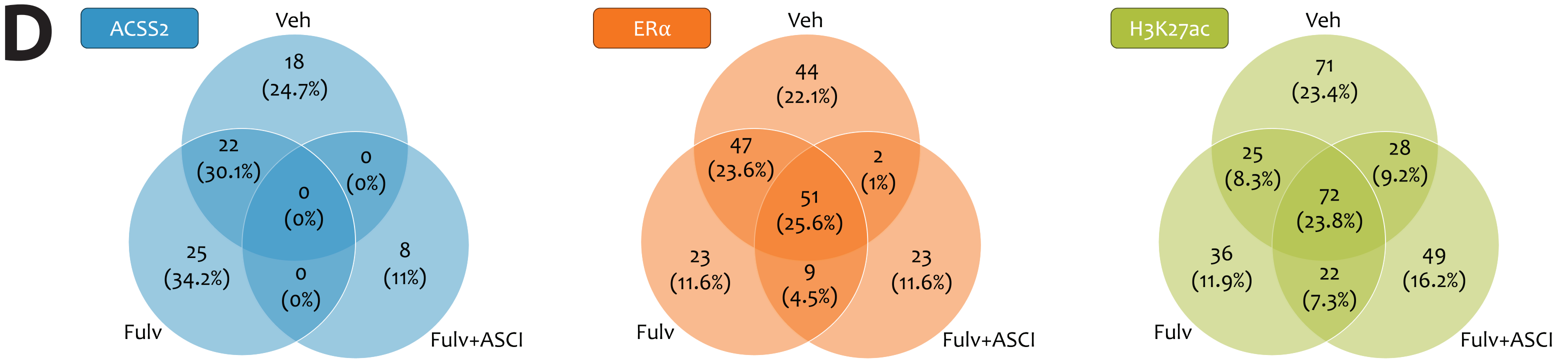


# E

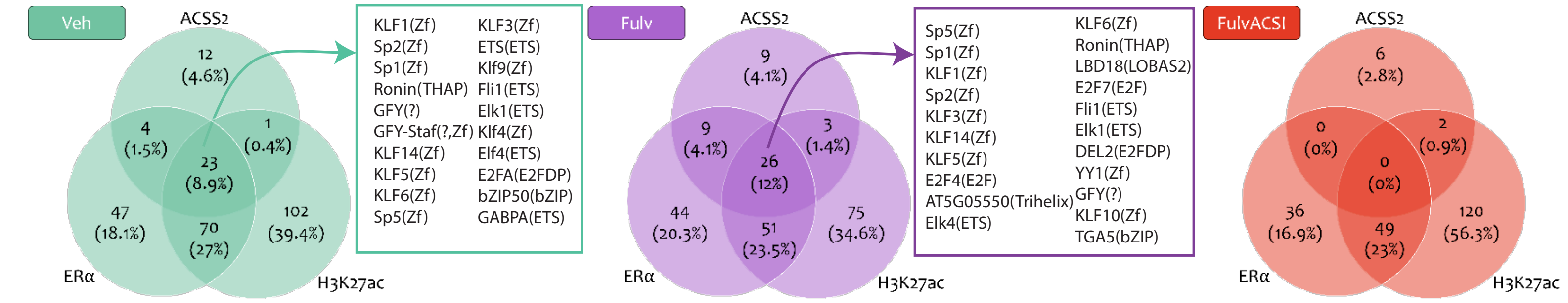


### Supplementary Figure 3
