## Supplemental Figure 6 for "ACSS2-Mediated Metabolic-Epigenetic Crosstalk Drives Fulvestrant Resistance and Represents a Novel Therapeutic Target"

#### Homer Known Motif Enrichment Results

| Rank | Motif | Name | P-value | % of Targets Sequences with Motif | % of Background Sequences with Motif |
|---|---|---|---|---|---|
| 1 | 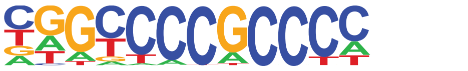 | Sp2(Zf)/HEK293-Sp2.eGFP-ChIP-Seq(Encode)/Homer | 1e-28 | 18.89% | 7.93% |
| 2 | 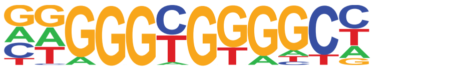 | KLF1(Zf)/HUDEP2-KLF1-CutnRun(GSE136251)/Homer | 1e-24 | 12.43% | 4.36% |
| 3 | 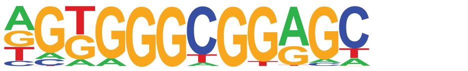 | Sp5(Zf)/mES-Sp5.Flag-ChIP-Seq(GSE72989)/Homer | 1e-21 | 13.32% | 5.31% |
| 4 | 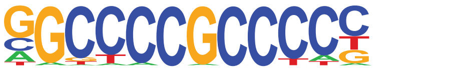 | Sp1(Zf)/Promoter/Homer | 1e-21 | 7.55% | 2.03% |
| 5 | 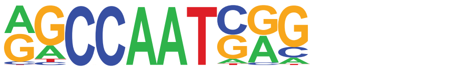 | NFY(CCAAT)/Promoter/Homer | 1e-20 | 5.77% | 1.22% |
| 6 | 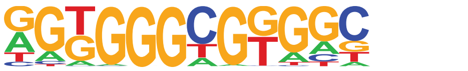 | KLF14(Zf)/HEK293-KLF14.GFP-ChIP-Seq(GSE58341)/Homer | 1e-17 | 18.09% | 9.27% |
| 7 | 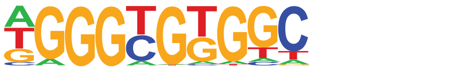 | KLF5(Zf)/LoVo-KLF5-ChIP-Seq(GSE49402)/Homer | 1e-13 | 13.02% | 6.50% |
| 8 | 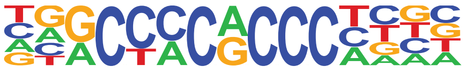 | KLF3(Zf)/MEF-Klf3-ChIP-Seq(GSE44748)/Homer | 1e-11 | 5.96% | 2.10% |
| 9 | 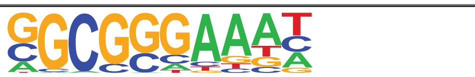 | E2F4(E2F)/K562-E2F4-ChIP-Seq(GSE31477)/Homer | 1e-9 | 5.86% | 2.39% |
| 10 | 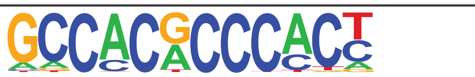 | Klf9(Zf)/GBM-Klf9-ChIP-Seq(GSE62211)/Homer | 1e-7 | 4.27% | 1.63% |
| 11 | 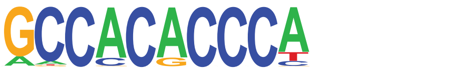 | Klf4(Zf)/mES-Klf4-ChIP-Seq(GSE11431)/Homer | 1e-7 | 3.38% | 1.15% |
| 12 | 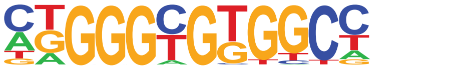 | KLF6(Zf)/PDAC-KLF6-ChIP-Seq(GSE64557)/Homer | 1e-6 | 9.15% | 5.14% |

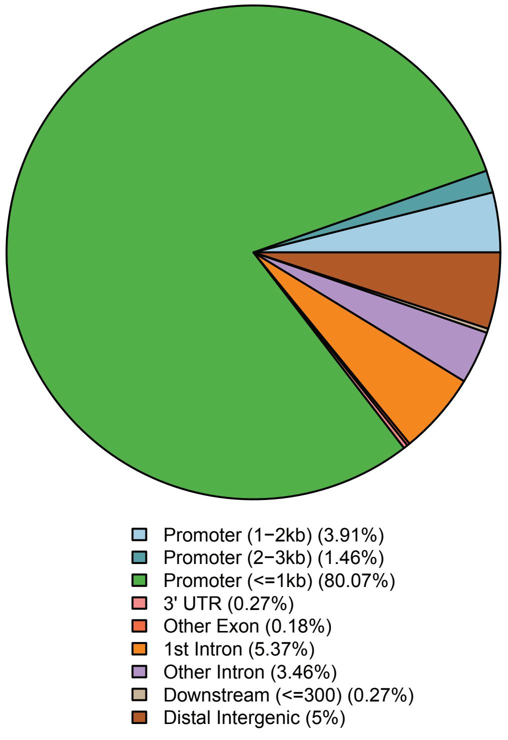


### Supplementary Figure 6
