## Supplemental Table 1 for "ACSS2-Mediated Metabolic-Epigenetic Crosstalk Drives Fulvestrant Resistance and Represents a Novel Therapeutic Target"

|  | Cluster 1 | Cluster 2 |
| --- | --- | --- |
| CP | - Aerobic glycolysis - augmented - Aerobic glycolysis - Metabolic reprogramming in colon cancer - Alanine, aspartate and glutamate metabolism - Warburg effect modulated by deubiquitinating enzymes and their substrates - Glycolysis and gluconeogenesis - Metabolic reprogramming in pancreatic cancer - Clear cell renal cell carcinoma pathways - Glycolysis / Gluconeogenesis - Metabolic epileptic disorders - Amino acid metabolism - Arginine and proline metabolism - Glucose metabolism - Metabolism of nucleotides - Cellular response to starvation - Aerobic respiration and respiratory electron transport - Metabolism of carbohydrates - Metabolism of amino acids and derivatives - Cellular responses to stimuli - Infectious disease | - NFE2L2 regulating anti-oxidant/detoxification enzymes - Pathway Definition from KEGG: (O2-,HO2,H2O2,OH,ACRL,4HNE,NO) -\| KEAP1 -\| NRF2 => (HMOX1,NQO1,GST,TXNRD1) - Oxidative stress response - Nuclear events mediated by NFE2L2 - KEAP1-NFE2L2 pathway - Clock-controlled autophagy in bone metabolism - TP53 Regulates Metabolic Genes - Cellular response to chemical stress - Androgen receptor signaling - DNA damage response (only ATM dependent) - Corticotropin-releasing hormone signaling - Androgen receptor network in prostate cancer - NRF2 pathway - Chromatin modifying enzymes - Nuclear receptors meta-pathway - Cancer pathways - Pathways in cancer - Pleural mesothelioma - Cellular responses to stimuli - RNA Polymerase II Transcription |
| GO | - The chemical reactions and pathways involving an L-amino acid. [GOC:edw] - The chemical reactions and pathways involving an alpha-amino acid. [GOC:TermGenie] - The chemical reactions and pathways involving glucose, the aldohexose gluco-hexose. D-glucose is dextrorotatory and is sometimes known as dextrose; it is an important source of energy for living organisms and is found free as well as combined in homo- and hetero-oligosaccharides and polysaccharides. [ISBN:0198506732] - The chemical reactions and pathways resulting in the formation of ribose phosphate, any phosphorylated ribose sugar. [GOC:ai] - The chemical reactions and pathways involving a nucleoside triphosphate, a compound consisting of a nucleobase linked to a deoxyribose or ribose sugar esterified with triphosphate on the sugar. [GOC:go_curators, ISBN:0198506732] - The chemical reactions and pathways involving monosaccharides, the simplest carbohydrates. They are polyhydric 2alcohols containing either an aldehyde or a keto group and between three to ten or more carbon atoms. They form the constitutional repeating units of oligo- and polysaccharides. [ISBN:0198506732] - The chemical reactions and pathways involving amino acids, carboxylic acids containing one or more amino groups. [ISBN:0198506732] - The chemical reactions and pathways resulting in the formation of a nucleoside phosphate. [GOC:TermGenie] - The chemical reactions and pathways involving ribose phosphate, any phosphorylated ribose sugar. [GOC:ai] - The chemical reactions and pathways resulting in the breakdown of small molecules, any low molecular weight, monomeric, non-encoded molecule. [GOC:curators, GOC:vw] - The chemical reactions and pathways resulting in the formation of precursor metabolites, substances from which energy is derived, and any process involved in the liberation of energy from these substances. [GOC:jl] - The chemical reactions and pathways involving a purine-containing compound, i.e. any compound that contains purine or a formal derivative thereof. [GOC:mah] - The cellular chemical reactions and pathways involving a nucleobase-containing small molecule: a nucleobase, a nucleoside, or a nucleotide. [GOC:vw] - The chemical reactions and pathways involving organic acids, any acidic compound containing carbon in covalent linkage. [ISBN:0198506732] - The chemical reactions and pathways involving carbohydrates, any of a group of organic compounds based of the general formula Cx(H2O)y. [GOC:mah, ISBN:0198506732] - The chemical reactions and pathways resulting in the formation of small molecules, any low molecular weight, monomeric, non-encoded molecule. [GOC:curators, GOC:pde, GOC:vw] - The chemical reactions and pathways involving organophosphates, any phosphate-containing organic compound. [ISBN:0198506732] - The chemical reactions and pathways involving small molecules, any low molecular weight, monomeric, non-encoded molecule. [GOC:curators, GOC:pde, GOC:vw] - The chemical reactions and pathways involving carbohydrate derivative. [GOC:TermGenie] - The chemical reactions and pathways resulting in the formation of organonitrogen compound. [GOC:pr, GOC:TermGenie] | - The chemical reactions and pathways involving a pyridine-containing compound, i.e. any compound that contains pyridine or a formal derivative thereof. [GOC:mah] - Any process that results in a change in state or activity of a cell or an organism (in terms of movement, secretion, enzyme production, gene expression, etc.) as a result of a toxic stimulus. [GOC:lr] - The chemical reactions and pathways resulting in the formation of organic acids, any acidic compound containing carbon in covalent linkage. [ISBN:0198506732] - Any process that results in a change in state or activity of a cell or an organism (in terms of movement, secretion, enzyme production, gene expression, etc.) as a result of a stimulus reflecting the presence, absence, or concentration of nutrients. [GOC:mah] - The process whose specific outcome is the progression of a gland over time, from its formation to the mature structure. A gland is an organ specialised for secretion. [GOC:jid] - The chemical reactions and pathways involving a purine-containing compound, i.e. any compound that contains purine or a formal derivative thereof. [GOC:mah] - The chemical reactions and pathways resulting in the formation of small molecules, any low molecular weight, monomeric, non-encoded molecule. [GOC:curators, GOC:pde, GOC:vw] - A cellular process involving delivery of a portion of the cytoplasm to lysosomes or to the plant or fungal vacuole that does not involve direct transport through the endocytic or vacuolar protein sorting (Vps) pathways. This process typically leads to degradation of the cargo; however, it can also be used to deliver resident proteins, such as in the cytoplasm-to-vacuole targeting (Cvt) pathway. [PMID:21997368, PMID:22966490, PMID:28596378] - Catalysis of an oxidation-reduction (redox) reaction, a reversible chemical reaction in which the oxidation state of an atom or atoms within a molecule is altered. One substrate acts as a hydrogen or electron donor and becomes oxidized, while the other acts as hydrogen or electron acceptor and becomes reduced. [GOC:go_curators] - The cellular chemical reactions and pathways involving a nucleobase-containing small molecule: a nucleobase, a nucleoside, or a nucleotide. [GOC:vw] - The chemical reactions and pathways involving organic acids, any acidic compound containing carbon in covalent linkage. [ISBN:0198506732] - The chemical reactions and pathways resulting in the breakdown of substances, carried out by individual cells. [GOC:jl] - Any process that results in a change in state or activity of a cell or an organism (in terms of movement, secretion, enzyme production, gene expression, etc.) as a result of a lipid stimulus. [GOC:sl] - The chemical reactions and pathways involving organophosphates, any phosphate-containing organic compound. [ISBN:0198506732] - The chemical reactions and pathways involving small molecules, any low molecular weight, monomeric, non-encoded molecule. [GOC:curators, GOC:pde, GOC:vw] - Any process that modulates the frequency, rate or extent of cell differentiation, the process in which relatively unspecialized cells acquire specialized structural and functional features. [GOC:go_curators] - Any process that results in a change in state or activity of a cell or an organism (in terms of movement, secretion, enzyme production, gene expression, etc.) as a result of an oxygen-containing compound stimulus. [GOC:pr, GOC:TermGenie] - Any process that activates or increases the rate or extent of development, the biological process whose specific outcome is the progression of an organism over time from an initial condition (e.g. a zygote, or a young adult) to a later condition (e.g. a multicellular animal or an aged adult). [GOC:ai] - The chemical reactions and pathways involving lipids, compounds soluble in an organic solvent but not, or sparingly, in an aqueous solvent. Includes fatty acids; neutral fats, other fatty-acid esters, and soaps; long-chain (fatty) alcohols and waxes; sphingoids and other long-chain bases; glycolipids, phospholipids and sphingolipids; and carotenes, polyprenols, sterols, terpenes and other isoprenoids. [GOC:ma] - Any process that results in a change in state or activity of a cell (in terms of movement, secretion, enzyme production, gene expression, etc.) as a result of a stimulus indicating the organism is under stress. The stress is usually, but not necessarily, exogenous (e.g. temperature, humidity, ionizing radiation). [GOC:mah] |
